# Resource Quality Differentially Impacts *Daphnia* Interactions with Two Parasites

**DOI:** 10.1101/2024.07.31.605800

**Authors:** Michelle L. Fearon, Kristel F. Sánchez, Syuan-Jyun Sun, Siobhan K. Calhoun, Kira J. Monell, Varun Ravichandran, Meghan A. Duffy

## Abstract

Resource quality can have conflicting effects on host–parasite interactions; for example, higher resource quality might increase host investment in immune function, or conversely, might permit greater parasite reproduction. Thus, anticipating the impact of changing resource quality on host– parasite interactions is challenging, especially because we often lack a mechanistic understanding of how resource quality influences host physiology and fitness to alter infection outcomes. We investigated whether there are generalizations in how resource quality affects multiple host clones’ interactions with different parasites. We used the *Daphnia* freshwater zooplankton model system to experimentally investigate how a resource quality gradient from high-quality green algae to poor-quality cyanobacteria diets influences host fitness, physiology, and infection by two parasites: a bacterium, *Pasteuria ramosa*, and a fungus, *Metschnikowia bicuspidata*. We ran a separate experiment for each parasite using a factorial design with four diets, two *Daphnia dentifera* host clones, and parasite-inoculated and uninoculated treatments (16 treatments per experiment). Diet strongly influenced infection by the fungus but not the bacterium. These relationships between diet and infection cannot be explained by changes in feeding rate (and, therefore, parasite exposure). Instead, the impact of diet on fungal infection was associated with impacts of diet on the earliest stage of infection: hosts that fed on poor quality diets had very few attacking spores in their guts. Diet did not significantly influence host immune responses. Diet influenced spore production differently for the two parasites, with reduced resource quality limiting the number of fungal spores and the size (but not number) of bacterial spores. Diet, host clone, and infection all affected host fitness. Interestingly, diet influenced the impact of the bacterium, a parasitic castrator that induces gigantism; for one clone, infected hosts fed high quality diets still produced a substantial number of offspring, whereas resource limitation hindered gigantism. Finally, there were often costs of resisting infection, though these generally were not affected by diet. Overall, we show that resource quality differentially impacts the exposure, infection, and proliferation processes for different parasites and host clones, which highlights the need to use multi-genotype and multi-parasite studies to better understand these complex interactions.

## Introduction

Anthropogenic effects on the environment, such as habitat loss and degradation, often alter the abundance and quality of food resources (Robb et al. 2008, Oro et al. 2013, Civitello et al. 2018). These changes in resources can impact disease dynamics in complicated ways. Reduced resource quality can limit host densities, reducing the pool of susceptible hosts for parasites to infect, and can alter host investment in immune function, reproduction, and survival (Beldomenico et al. 2008, Becker and Hall 2014). Resource quality can also influence the host’s exposure rate to environmentally transmitted parasites and a parasite’s ability to produce sufficient infectious propagules (Penczykowski et al. 2014, Budischak and Cressler 2018). Therefore, the consequences of anthropogenic alterations in resource quality on host–parasite interactions will depend on the sensitivity of host and parasite traits to changing resource quality in the environment (Mugabo et al. 2019). However, because we lack a mechanistic understanding of how resource quality alters host physiology, we are currently unable to predict how changing resource landscapes will alter how hosts interact with their parasites.

Resource quality can have conflicting effects on host–parasite interactions at each step of the infection process, from parasite exposure to initiation of infection to parasite reproduction. Many hosts shift their foraging behavior over time and space in response to resource quality (e.g., Mella et al. 2018, Weterings et al. 2018, Iannino et al. 2023), which could change the host’s exposure rate to environmental parasites. For example, hosts may increase foraging to take advantage of a high-quality resource (e.g., Schatz and McCauley 2007) or, alternatively, the host may increase foraging in response to a poor-quality resource to meet nutritional needs (e.g., Heuermann et al. 2011); in the (common) case where hosts encounter the parasite while foraging, these changes should increase the rate of parasite encounter. Upon initiation of the infection process, resource quality may influence the host’s ability to up-regulate energetically costly physiological or immunological defenses to adequately respond to the infection (Sandland and Minchella 2003, Cressler et al. 2014a, Cotter and Al Shareefi 2022). Additionally, in some cases, a secondary metabolite that renders a food low quality also provides protection against infection (e.g., Huffman and Caton 2001, Sánchez et al. 2024). Finally, once an infection is fully established, resource quality influences both the parasite and host. For instance, higher quality resources may allow the parasite to increase its growth and replication within the host (Hall et al. 2009). Alternatively, high-quality resources may allow the host to increase investment in immune function, limiting parasite production (e.g., Cornet et al. 2013).

Intraspecific variation in host responses further increases the challenge of predicting how changing resources will influence the outcomes of host–parasite interactions. There is often substantial variation within species in susceptibility to infection; host genotype often strongly influences susceptibility due to a variety of factors, including variation in cellular immunity and barrier defenses (e.g., Leitão et al. 2019, Cáceres and Stewart Merrill 2023). Similarly, susceptible host genotypes can vary in their quality for the parasite, with parasites able to produce more infectious propagules in certain host genotypes (e.g., Auld et al. 2017, Kutzer et al. 2023). Additionally, host genotypes can vary in their sensitivity to changing resource quality, with variation in tolerance of low resource quality, and, conversely, in ability to exploit rich resource environments (e.g., Watt 1986, Hall et al. 2012). These differences in how host genotypes allocate available resources to life history traits, such as reproduction, growth, and survival, along resource gradients and in the presence or absence of parasites should influence outcomes for both the host and the parasite. Moreover, host species are infected by a multitude of parasites, which may differ in how they are influenced by changing resource conditions.

Here, we experimentally manipulated a planktonic host-parasite system with a goal of understanding whether it is possible to predict the impacts of altered food quality on host-parasite interactions based on changes in host physiology and life history traits. To explore the generality of these patterns, we used two genotypes of the host (the freshwater zooplankton grazer, *Daphnia dentifera*) and two parasites (the fungus *Metschnikowia bicuspidata* and the bacterium *Pasteuria ramosa*), both of which are incidentally ingested while foraging on phytoplankton. We particularly focused on shifts in food quality related to the amount of the cyanobacterium *Microcystis aeruginosa* in the diet. Blooms of cyanobacteria, including *Microcystis*, are increasingly common as a result of nutrient pollution and climate change (Carpenter 2008, Huisman et al. 2018, Smucker et al. 2021). *Microcystis* is a low quality food for *Daphnia* (Agrawal et al. 2005, Schwarzenberger et al. 2012) but also can protect *Daphnia* from fungal infections (Sánchez et al. 2019, Manzi et al. 2020). Therefore, it is hard to predict the net impact of increased *Microcystis* on host–parasite interactions, and those changes may not be consistent across parasites. We conducted a factorial experiment that included four resource quality diets along a gradient, two host genotypes (hereafter, clones), and two parasite inoculation treatments (inoculated vs. uninoculated), yielding a total of 16 treatments for each of the two parasites. In general, we expected that reducing resource quality would reduce host growth rate, immune response, metabolic rate, reproductive rate, and survival, as well as reduce parasite exposure (through lower feeding rates), infection, and infectious propagule yield. We also expected that the two host clones would exhibit overall similar patterns but with slightly different sensitivities to changing diet and exposure to parasites; we expected that these differences would reflect variation in host resource acquisition and allocation of resources to different aspects of host life history. Similarly, since the parasites share many key traits, such as infection via foraging and exploitation of within host resources for reproduction, we predicted that reducing resource quality would reduce both parasites’ infectivity and yield. Overall, our goal was to untangle how reducing resource quality changes outcomes of parasitism via altered resource allocation, with an aim of improving our understanding of how environmental change will influence host–parasite interactions. By studying two host clones and two parasite species, we sought to assess the generality of responses of hosts and parasites to changing resource quality.

## Methods

### Host–Parasite System

*Daphnia dentifera* is a zooplankton species commonly found in small, stratified freshwater lakes in the Midwestern USA (Tessier and Woodruff 2002). Two common parasites of *D. dentifera* are *Metschnikowia bicuspidata* and *Pasteuria ramosa* (hereafter, the ‘fungus’ and ‘bacterium’, respectively), which hosts encounter while filter feeding on phytoplankton in the water column (Ebert 2005). Both parasites’ transmission spores penetrate through the host gut wall and proliferate using the host’s resources within the hemolymph, ultimately killing the host and releasing spores back into the water column (Ebert et al. 2015, Stewart Merrill and Cáceres 2018). While the fungus and bacterium share many key traits, they differ in some important ways. The bacterium has strong host–parasite genotype interactions, where infection success is dependent on compatible host and parasite genotypes (Carius et al. 2001, Luijckx et al. 2011, Bento et al. 2017). In contrast, infection by the fungus does not show host–parasite genotype specificity (Duffy and Sivars-Becker 2007, Auld et al. 2012b, Searle et al. 2015). Additionally, the two parasites differ in their primary fitness effects; the fungus increases mortality at a younger age, while the bacterium castrates the host, strongly reducing fecundity (Ebert et al. 2000, Auld et al. 2012b, Auld et al. 2017).

### Experimental Design

We conducted two sequential experiments that factorially manipulated resource quality, host genotype, and parasite exposure (4 diet treatments × 2 host genotypes × 2 parasite exposures (i.e., inoculated or uninoculated) = 16 treatment combinations per experiment). The first experiment used the bacterium in the parasite-inoculated treatment, while the second used the fungus for inoculation. We ran separate experiments for each parasite due to the number of treatments and response variables measured. Each treatment had 10 replicate animals in individual 50 mL beakers (N = 160 animals per experiment as the baseline, but with 80 additional animals in the fungus experiment – described below – for a total of 240 animals in the fungus experiment). We conducted the bacterium experiment from 9 Nov–16 Dec 2021, and the fungus experiment from 4–27 Jan 2022. The resource quality gradient varied from 100% high-quality green algae, *Scenedesmus obliquus* (UTEX 3155) to 100% low-quality blue-green cyanobacteria, *Microcystis aeruginosa* (NIVA CYA43), with a 50:50 mixture of the two to achieve an intermediate resource quality. Both phytoplankton strains are single-celled and edible for *Daphnia* but vary in their nutritional quality (as confirmed in this study). The *Scenedesmus* and *Microcystis* cultures were grown in chemostats with COMBO and Z8 media, respectively, under 24 h light at room temperature. The phytoplankton food for each diet was prepared weekly for both experiments by centrifuging the phytoplankton, pouring off the supernatant, and resuspending the food in Artificial *Daphnia* Medium (ADaM; Klüttgen et al. 1994); see below for more information.

*Microcystis* strain CYA43 does not produce microcystin toxin (Burberg et al. 2020 and confirmed in this experiment (see below)), but because microcystin toxin is commonly produced in harmful *Microcystis* blooms in lakes (Chaffin et al. 2023), we also included a toxin-spiked treatment with 15 µg/L microcystin-LR (Cayman Chemical Company, Ann Arbor, MI). This represents a high but ecologically relevant level of microcystin-LR that is within the range observed during natural *Microcystis* blooms (Ibelings et al. 2005, Ha et al. 2009, Hayes et al. 2020, Chaffin et al. 2023) and is less than the 24 µg/L guideline value for recreational waters (Chorus and Welker 2021). Therefore, our four resource quality treatments (that is, diets) were 100% *Scenedesmus* (S), 50:50 *Scenedesmus*:*Microcystis* (SM), 100% *Microcystis* (M), and 100% *Microcystis* + microcystin toxin (M+). The M and M+ treatments represent the same low resource quality, so the comparison between these two treatments allowed us to evaluate the effects of the toxin on host physiology and infection outcomes.

We collected *D. dentifera* neonates (<24 hours old) from the ‘Mid37’ clone (originally from Midland Lake, Greene County, Indiana, USA) and ‘Standard’ clone (originally isolated from Barry County, MI, USA). To do this, we maintained stock cultures of both the Mid37 clone and Standard clone for at least six generations to standardize maternal effects. The stock cultures were fed 2 mg dry weight/L *Scenedesmus* three times per week; approximately 50% of the dry weight of phytoplankton comes from carbon (Grobbelaar 2004, Huang et al. 2018). The stock cultures and experimental animals were maintained at 20°C and 16:8 light:dark cycle throughout the entire experiment. Both experiments followed the same procedure. On Day 1 of the experiment, we moved host individuals to new beakers filled with filtered lake water. We collected over 100 neonates (<24 h old) on Day 2 and placed 10 animals per beaker in 200 mL filtered lake water; each beaker was fed 2 mg dry weight/L *Scenedesmus* food and maintained for two days. On Day 4, we moved animals to individual 50 mL beakers with 30 mL of fresh filtered lake water and haphazardly assigned individuals to diet treatments. Animals were maintained on their new diet treatment for 2 days. Throughout both experiments, animals were fed the same concentration of food in their beakers during each feeding, unless otherwise noted: the S diet treatment was given 2 mg dry weight/L *Scenedesmus*, the SM diet treatment received 1 mg dry weight/L *Scenedesmus* and 1 mg dry weight/L *Microcystis*, and the M and M+ diet treatments were given 2 mg dry weight /L *Microcystis*. For the M+ treatment, we also added 15 µg/L microcystin-LR each time the animals were fed. We confirmed microcystin concentrations in our beakers prior to and during parasite exposure with ELISAs (Gold Standard Diagnostics Horsham, Inc., Warminster, PA) following the manufacturer’s protocol; M+ treatments had 16.45 ± 2.15 µg/L (mean ± SE) and 17.93 ± 1.36 µg/L microcystin concentrations on average in the fungus and bacterium experiments, respectively, while all other diets had 0.17 ± 0.017 µg/L and 0.20 ± 0.019 µg/L in the two experiments from the filtered lake water (Figure S1). In the bacterium experiment on Day 4, the M+ treatment accidentally received 150 µg/L microcystin for one day due to a calculation error, before we corrected the mistake by moving those animals to new beakers with filtered lake water and fresh M+ food and 15 µg/L microcystin-LR. We confirmed the toxin “super spike” with ELISAs (Figure S2). Only a single replicate died during this super spike (M+, Mid37, *Pasteuria*-inoculated, Rep 4) but it did not appear to otherwise affect the experiment.

On Day 6, we added either 2,000 bacterial spores/mL (isolate G/18, originally isolated from Midland Lake, Greene County, Indiana, USA; experiment 1) or 500 fungal spores/mL (“Standard” isolate, originally from Baker Lake, Barry County, MI, USA; experiment 2) to the parasite-inoculated treatments. Prior work has shown that a lower dose of the fungus achieves similar levels of infection as the higher dose of the bacterium (Auld et al. 2012b, 2014). An equivalent volume of ground *Daphnia* carcasses was added to the uninoculated treatments as a sham-exposure. All treatments received a half quantity (1 mg dry weight/L total) of their assigned diet during exposure and M+ treatments also received 15 µg/L microcystin. On Day 8, after 48 hours, we moved all animals to new beakers with spore-free filtered lake water and added the full 2 mg dry weight/L of each diet. For the remainder of both experiments, all animals received two water changes per week, were checked for offspring and mortality at least four times per week, and were given five additions of 2 mg/L food per week according to their diet treatment and maintained in incubators at 20°C and 16:8 light:dark cycle. We measured feeding rate and respiration rate for each animal during parasite exposure (described in more detail in the <u>Parasite and Host Measurements section</u> below and Appendix S1: Section S1). The bacterium experiment was concluded on Day 38 (30 days post-inoculation), and the fungus experiment was concluded on Day 24 (16 days post-inoculation). The fungal infections develop more rapidly, causing host mortality sooner than bacterial infections (Ebert et al. 2000, Auld et al. 2014). At the conclusion of the experiments, we assessed the remaining animals for infection under a dissecting microscope at 40-60x magnification and measured their body length (detailed methods described in Appendix S1: Section S1), before storing them at -20°C in individual 1.5 mL microcentrifuge tubes with 0.1 mL deionized water for spore counts.

In the fungus experiment, we also included an additional set of 10 replicate animals for each diet × clone treatment that was inoculated with the fungus (yielding the 80 additional animals mentioned above). These animals allowed us to quantify parasite exposure in their guts and hemocyte immune response one day post-inoculation (Day 9) following established methods (Stewart Merrill et al. 2019, Sun et al. 2023). These replicates were in addition to the baseline experimental animals in case the combined effects of parasite inoculation and quantification of spores in host guts was stressful and decreased survival. The Standard SM, M, and M+ diet treatments only had 9 additional replicates instead of 10 because there were insufficient fungal spores to infect these replicates; the extra 10^th^ replicate for the SM, M, and M+ diets were marked as ‘uninoculated’ and maintained for the duration of the experiment. We counted and summed the total number of attacking fungal spores that were embedded in or penetrated the gut wall or were entirely in the haemocoel as a measure of fungal parasite exposure (Stewart Merrill et al. 2019). To evaluate the cellular immune response, we counted the total number of hemocytes that were attached to fungal spores that had entered the hemocoel (i.e., penetrated or hemocoel spores). We also calculated the average number of hemocytes per spore by dividing the total number of hemocytes by the number of penetrated and hemocoel spores. Animals that survived the procedure were then maintained through the end of the experiment under the same conditions as all other treatments (described above), and we collected offspring production, infection, and parasite yield data (described in detail below). Animals that died due to the procedure to quantify parasite exposure in their guts ranged from 1 – 7 per diet × host clone combination (mean = 4.1), with greater mortality in the S and SM treatments (5 – 7 per diet x clone combination, mean = 6) compared to the M and M+ treatments (1 – 4 per diet x clone combination, mean = 2.25).

We were unable to evaluate within host bacterial parasite exposure and immune responses in the same way. Attachment of the bacterium endospore to the esophagus is determined by genotypic specificity, and we already knew that this bacterial isolate could infect these host genotypes. In contrast to the fungus, it is unknown what happens in the early stages of infection by the bacterium (Ebert et al. 2015). Moreover, prior work suggests that immune responses are symptomatic of bacterial infection but do not mediate recovery (Auld et al. 2012a).

### Parasite and Host Measurements

We quantified offspring production, survival, infection status, attacking spores (fungus only), mature spore yield, spore size (bacterium only), feeding rate, metabolic rate, and host body size and growth rate using established methods. More detail is provided in Appendix S1. Methods for attacking spores (Stewart Merrill et al. 2019), feeding rate (Hite et al. 2020), and metabolic rate (Nørgaard et al. 2021) were based on the literature.

### Statistical Analyses

All analyses were run in R version 4.2.1 (R Core Team 2022). We analyzed the data from the fungus and bacterium experiments separately since the two experiments were run independently. For all generalized linear models, we tested for overdispersion and zero-inflation and selected the best model distributions using the model with the lowest AIC and the best fit of the diagnostic tests (DHARMa package; Hartig 2020). For all linear regressions, we checked for normality of the residuals and model assumptions using visual diagnostic residual plots. We then evaluated overall effects of each main factor with a Type II Analysis of Deviance test (car package; Fox and Weisberg 2019) and a Tukey-adjusted post-hoc test to determine the differences between diet treatments for each host clone (emmeans package; Lenth 2022).

#### Parasite infectivity and host susceptibility

For both parasites, we evaluated how infection prevalence varied based on host clone and diet, and the interaction between those factors. We defined “infection” as animals that contained mature transmission spores at death or the end of the experiment; this definition is analogous to the “terminal infection” designation of Stewart Merrill et al. (2019). Only one animal in the fungus experiment and one in the bacterium experiment had immature spores but no mature spores at death. Using animals that were inoculated with parasites, we modeled infection prevalence with a generalized linear mixed-effects model (GLMM) with a binomial distribution and logit link function (lme4 package; Bates et al. 2015) and block as a random effect (each experiment had two blocks, which determined the order of feeding and respiration rate measurements, see Appendix S1: Section S1). Animals that died prior to experiment day 14, which was 7 days after inoculation, were excluded from the analysis of infection prevalence because symptoms of infection were not apparent before this point (excluded: nfungus = 64, nbacterium = 25); the relatively high number of animals that died early in the fungus experiment was a result of checking for attacking spores in the additional animals (as described above). In the fungus infection prevalence model, because the M and M+ treatments in the Mid37 clone failed to produce infections, we removed those treatments and the non-significant interaction term from the analysis because the lack of variation in those treatments led to separation in the data and very large standard errors in the model. However, we still included the Mid37 clone M and M+ treatments in Figure 1a to demonstrate the lack of infection. Infection prevalence and confidence intervals were calculated using the epiR package (Stevenson et al. 2020).

**Figure 1.**
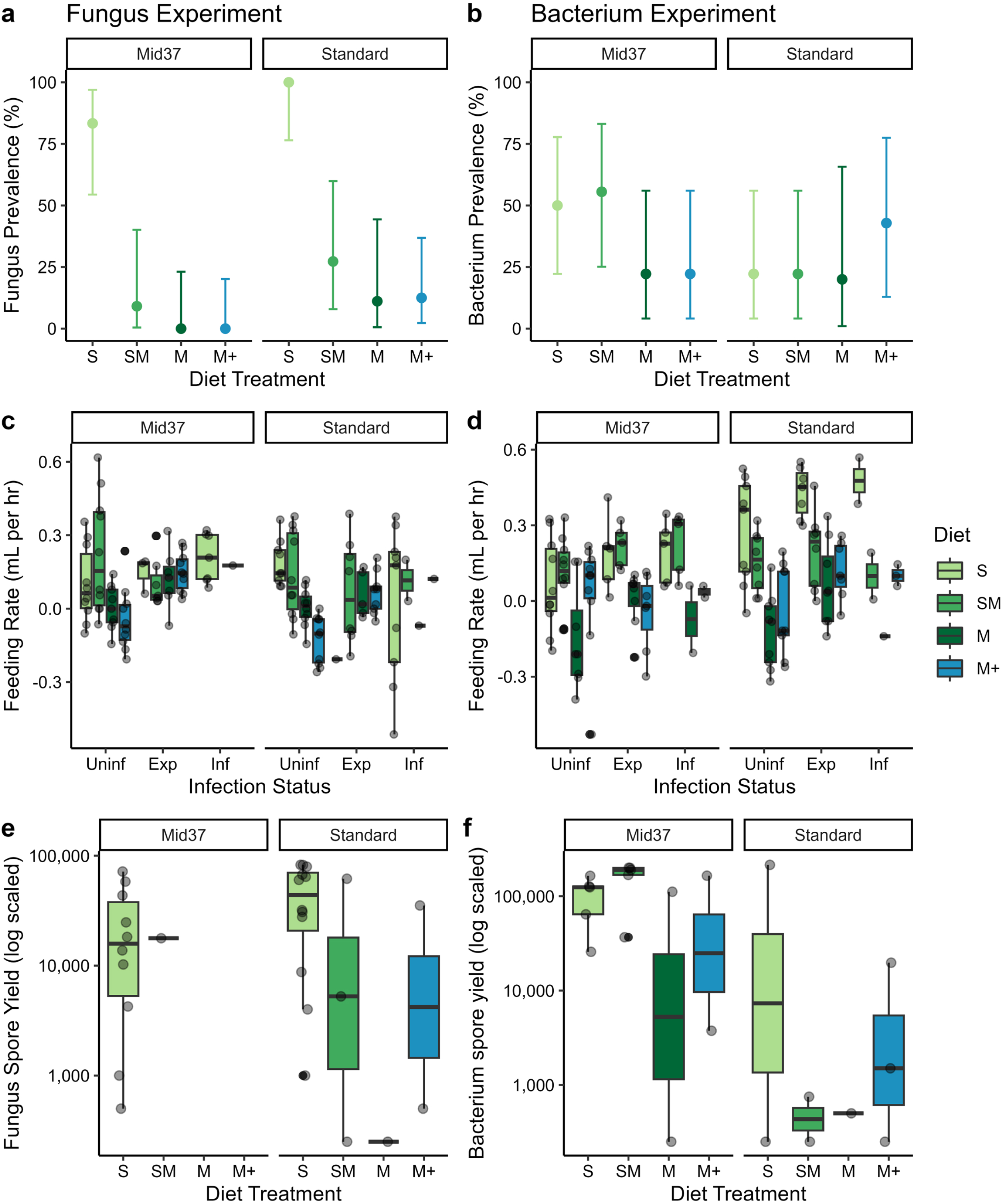
Reducing resource quality protected hosts from fungal infection (a), reduced host feeding rate in uninfected hosts but not fungus exposed or infected hosts (c), and reduced fungus mature spore yield (e). Resource quality did not impact bacterial infection prevalence (b) nor the mature spore yield (f), but did alter host feeding rates (d). a,b) Fungal and bacterial terminal infection prevalence with 95% confidence intervals for each resource quality diet and host clone treatment combination. c,d) Individual host filter feeding rate (mL per hour) – a proxy for parasite exposure rate for parasite inoculated treatments – for each resource quality diet, host clone, and infection status combination. Infection status: Uninf = Uninfected (and uninoculated), Exp = Exposed-but-uninfected, Inf = Infected. e,f) Individual fungal and bacterial mature spore yield from infected hosts for each resource quality diet and host clone treatment combination. Fungal spore yield decreased with decreasing resource quality. Bacterial spore yield differed based on host clone but not diet; Standards had lower mature bacterial spore yield compared to Mid37 clones.

#### Parasite exposure and feeding rate

We evaluated the feeding rate during the parasite exposure period to assess the differences in energetic intake and parasite exposure based on diet, host clone, and infection status at the end of the experiment: uninoculated and uninfected (referred to as ‘uninfected’ below, though sometimes referred to with the longer ‘uninoculated and uninfected’ to reduce potential confusion), parasite inoculated but uninfected (hereafter ‘exposed-but-uninfected’), and parasite inoculated and infected (hereafter ‘infected’). We separated the parasite-inoculated animals that resisted infection from those that became infected to be able to test for differences in feeding behavior during parasite exposure that may have contributed to different infection outcomes.

Additionally, this approach allows us to test how host feeding behavior differed in the presence and absence of parasites by comparing the exposed-but-uninfected hosts to the uninfected hosts. We modeled feeding rate with a linear mixed effects model (LMM) that included diet, host clone, infection status, all two-way interactions, and block as a random effect (lme4 package; Bates et al. 2015). The bacterium experiment had sufficient data within each diet × clone × infection treatment combination to include the three-way interaction, but we were unable to include the three-way interaction in the model for the fungus experiment.

To further assess the differences between treatment groups, we conducted a series of Tukey-adjusted post-hoc tests (emmeans package; Lenth 2022). First, we evaluated the differences between diets conditional on host clone and infection status to assess how each host clone responded to decreasing resource quality. Second, we tested the contrasts between each infection status conditional on host clone and averaging across all the diets to be able to assess how each clone may have shifted their feeding behavior in response to the presence of parasite spores in the water.

#### Parasite attacking host gut and host hemocyte immune response (fungus only)

We tested the total number of fungal spores attacking the host gut based on diet, host clone, and host body length, and their interactions using a GLMM with a Poisson distribution and a log link function (glmmTMB package; Brooks et al. 2017). Host body length and its interaction terms were not significant in the model and were removed to improve the AIC of the remaining model with only diet, host clone, and diet × clone terms. These models only include the additional fungus-inoculated treatments that were assessed for spores in their gut and does not include the hosts that underwent the feeding and respiration assays.

For hosts with fungal spores that had partially or entirely punctured the host gut wall, we calculated the number of hemocyte immune cells per attacking fungus spore. We omitted embedded spores because these had not crossed the gut wall and therefore could not have hemocytes attached. We modeled hemocytes per spore using a GLMM with the number of hemocytes as the response variable, with the number of punctured spores as an offset and a Poisson distribution with a log link function. The model included diet and host clone as main factors. There were not enough replicates in all diet × clone treatments to include the interaction. We added an observation level random effect to the model to control for overdispersion in the initial model.

#### Mature parasite spore yield and size

For the animals that were infected, we modeled the production of bacterial and fungal mature spores per host (spore yield) with host clone, diet, and their interaction as main factors and block as a random effect in a GLMM with a quasipoisson distribution for the bacterium and a negative binomial distribution for the fungus, and a log link function for both models (glmmTMB package; Brooks et al. 2017). The fungal mature spore yield data had too few infected animals in all diet × clone treatments to include the interaction term in that model, but the bacterial mature spore yield model did include the interaction term. In additional models, we tested the effects of average microcystin concentration (both parasites) and number of hemocytes per spore (fungus only) on mature spore yield (methods and results in Appendix S1).

We used a linear regression to test for differences in bacterial spore size based on diet.

We were unable to adequately test for differences between the host clones because Standards had lower infection prevalence and produced fewer spores than Mid37s. We did not collect data on fungal spore size due to logistic constraints. However, prior work from our lab suggests that it is less likely that there would be a difference in fungal spore size based on resources: hosts coinfected with both the fungus and bacterium had smaller bacterial spores compared to singly infected hosts (presumably due to resource competition), but there was no difference in fungal spore size between coinfected and singly infected hosts (McLean 2022).

#### Host fecundity and survival

We evaluated how host total offspring production (total fecundity) varied based on diet, host clone, infection status, and all their interactions. We modeled total fecundity for each individual using a GLMM with a negative binomial distribution for the bacterium experiment and a quasipoisson distribution for the fungus experiment, and a log link function for both. The total fecundity data only included animals that survived beyond experimental day 11 (excluded: nfungus = 57 (23.8%), nbacterium = 23 (14.4%)), since that was the first day that we observed offspring in both experiments. The analysis for the bacterium experiment included all interactions including the three-way interaction between diet × clone × infection status, but the fungus model had insufficient power to assess the three-way interaction. We then conducted the same two sets of Tukey-adjusted post-hoc tests as previously described for the feeding rate analysis.

Finally, we assessed differences in host survival based on all treatment combinations using a Cox Proportional Hazards Ratio model to test for the risk of death (survival and survminer packages; Therneau and Grambsch 2000, Kassambara et al. 2011, Therneau 2022). We ran three versions of the hazard ratio models and produced a forest plot for each parasite experiment, including the full data set (all diet, clone, and infection status combinations), and one for each host clone (diet and parasite treatment data subset by clone). Since the hazard ratio models did not have sufficient power to allow for interaction terms, this approach allowed us to investigate the overall treatment effects and compare how diet and infection status affected risk of death for each host clone specifically. Animals that died early due to non-experimental causes (e.g., dropped beaker, animal handling, etc.) and the animals that lived to the end of the experiment were censored in the Cox Proportional Hazards Ratio model (Fungus: nnon-experimental cause = 1, nend of experiment = 139; Bacterium: nnon-experimental cause = 3, nend of experiment = 54). As with the fecundity analysis, we excluded animals that died on or prior to day 11 of the experiment (excluded: nfungus = 57 (23.8%), nbacterium = 23 (14.4%)). We used the likelihood ratio test to evaluate the overall importance of the variables included in the models, and a Wald test to evaluate the significance of each variable coefficient in the model.

For both the fecundity and survival analyses, 6 fungal-inoculated and 1 bacterium- inoculated animals that died on days 12-13, as well as 2 fungal-inoculated hosts that died on day 14, were coded as exposed-but-uninfected after spore counts did not find any spores.

#### Additional analyses (results in Supplement)

We conducted additional tests of the body size corrected feeding rate (i.e., relative feeding rate), host metabolic rate, host growth rate, and host body size during parasite exposure based on each treatment combination to explore the physiological drivers of differences in feeding rates (see Appendix S1: Section S1, Table S2).

## Results

### Parasite infectivity and host susceptibility

The effect of diet on infections differed for the two parasites, with strong effects of diet on infection by the fungus but not the bacterium (Figure 1a,b; Table 1). Decreasing resource quality reduced fungus infection prevalence such that hosts that consumed any of the poorer quality diets that included *Microcystis* (SM, M, and M+) were protected from fungus infection compared to the high-quality S diet (Figure 1a; χ^2^ diet = 18.0, df = 3, p < 0.001). Because the M and M+ diets for Mid37 failed to produce infections, we were unable to assess whether there was an interaction between diet and clone in this model. Conversely, bacterial infection prevalence was not significantly affected by changes in diet or host clone (Figure 1b, χ^2^ = 1.6, df = 3, p = 0.66; χ^2^ = 1.2, df = 3, p = 0.27), and there was no significant interaction between diet and clone.

**Table 1.** Type II analysis of deviance tables for each response variable for the fungus and bacterium experiments, respectively, presented in the main text: parasite terminal infection prevalence, host feeding rate (i.e., parasite exposure rate), fungal spores attacking the gut, hemocytes per attacking fungus spore, mature spore yield, and bacterial spore surface area. Significant p-values are bolded.

| Response Variable | Predictor Variables | Fungus Experiment |  |  | Bacterium Experiment |  |  |
| --- | --- | --- | --- | --- | --- | --- | --- |
|  |  | Chi-square | DF | P-value | Chi-square | DF | P-value |
| *Prevalence* | Diet | 18.02 | 3 | **0.0004** | 1.61 | 3 | 0.66 |
|  | Clone | 2.58 | 1 | 0.11 | 1.23 | 1 | 0.27 |
|  | Diet × Clone | -- | -- | -- | 3.33 | 3 | 0.34 |
| *Feeding Rate/ Parasite Exposure Rate* | Diet | 24.02 | 3 | **<0.0001** | 78.19 | 3 | **<0.0001** |
|  | Infection | 5.32 | 2 | 0.070 | 16.04 | 2 | **0.0003** |
|  | Clone | 6.73 | 1 | **0.0095** | 8.24 | 1 | **0.004** |
|  | Diet × Infection | 25.48 | 6 | **0.0003** | 4.83 | 6 | 0.57 |
|  | Diet × Clone | 1.80 | 3 | 0.61 | 12.91 | 3 | **0.005** |
|  | Infection × Clone | 5.67 | 2 | 0.059 | 1.33 | 2 | 0.51 |
|  | Diet × Infection × Clone | -- | -- | -- | 4.77 | 6 | 0.57 |
| *Fungal Spores Attacking Gut* | Diet | 183.51 | 3 | **<0.0001** | -- | -- | -- |
|  | Clone | 19.62 | 1 | **<0.0001** | -- | -- | -- |
|  | Diet × Clone | 1.53 | 3 | 0.68 | -- | -- | -- |
| *Hemocytes per Fungus Spore* | Diet | 2.63 | 3 | 0.45 | -- | -- | -- |
|  | Clone | 4.11 | 1 | **0.043** | -- | -- | -- |
|  | Diet × Clone | -- | -- | -- | -- | -- | -- |
| *Mature Spore Yield* | Diet | 19.85 | 3 | **0.0002** | 5.55 | 3 | 0.14 |
|  | Clone | 1.51 | 1 | 0.22 | 10.91 | 1 | **0.001** |
|  | Diet × Clone | -- | -- | -- | 2.55 | 3 | 0.47 |
| *Spore Surface Area* | Diet | -- | -- | -- | 3.64* | 3 | **0.039** |
| *Total Fecundity* | Diet | 63.06 | 3 | **<0.0001** | 85.25 | 3 | **<0.0001** |
|  | Infection | 49.59 | 2 | **<0.0001** | 26.44 | 2 | **<0.0001** |
|  | Clone | 160.03 | 1 | **<0.0001** | 50.89 | 1 | **<0.0001** |
|  | Diet × Infection | 4.27 | 6 | 0.64 | 2.48 | 6 | 0.87 |
|  | Diet × Clone | 30.07 | 3 | **<0.0001** | 9.76 | 3 | **0.021** |
|  | Infection × Clone | 6.37 | 2 | **0.041** | 12.80 | 2 | **0.002** |
|  | Diet × Infection × Clone | -- | -- | -- | 10.42 | 6 | 0.11 |

\* Estimate is an F value from an ANOVA.

### Parasite exposure and feeding rate

Host feeding rate during the inoculation period – and, consequently, the inadvertent parasite exposure rate – differed based on diet, host clone, and the host’s ultimate infection status (i.e., uninfected, exposed-but-uninfected, or infected) for both parasites (Figure 1c,d; Table 1). In the fungus experiment, the impact of decreasing resource quality on feeding rate differed based on the host’s ultimate infection status (Figure 1c; χ^2^ = 25.5, df = 6, p < 0.001). For uninoculated and uninfected hosts, reducing resource quality reduced feeding rate for both host clones. However, hosts that later became fungus-infected or that were fungus-exposed-but-uninfected did not differ in their feeding rates based on diet, indicating that animals in the parasite inoculation treatment consumed fungal spores at similar rates regardless of their diet (Table S3). For both clones, the feeding rates of fungus-exposed-but-uninfected and fungus-infected hosts were consistent (Table S4), indicating that a difference in feeding/exposure did not drive differences in whether hosts became infected.

In the bacterium experiment, host feeding rate also tended to decrease with reduced food quality, but the host clones differed somewhat in their responses to decreased resource quality (Figure 1d; χ^2^ = 12.9, df = 3, p = 0.005). Specifically, for Standards, feeding rate was highest for the high-quality S diet for all infection statuses. For Mid37s, hosts fed the SM diet had feeding rates similar to those fed the S diet, and higher than those fed the M diet (Table S5). In general, hosts that were bacterium-exposed-but-uninfected had higher feeding rates than uninoculated and uninfected hosts (χ^2^ = 16.0, df = 2, p < 0.001), contrary to expectations. Overall, exposed-but-uninfected Standards had feeding rates over three times higher than uninfected (and unexposed) Standards during the exposure period, while Mid37s with different ultimate infection (and inoculation) statuses did not significantly differ in their feeding behavior (Table S4).

Combining the results for infection prevalence and feeding rate during parasite inoculation (Figure 1a-d): in the fungus experiment, there was a strong effect of changing diet on infection prevalence but not on feeding rate of inoculated hosts. In the bacterium experiment, there was an effect of diet on feeding rate during inoculation but not on infection prevalence.

Moreover, in both experiments, the feeding rate during inoculation of hosts that ultimately became infected was not higher than the feeding rate of hosts that were exposed but did not become infected. Together, this suggests that, for both experiments, the patterns seen for infection prevalence were not driven by the rate of ingestion of spores.

### Parasite attacking host gut and host hemocyte immune response (fungus only)

Reducing resource quality strongly reduced the total number of embedded and penetrated fungal spores in the host gut wall (Figure 2a, χ^2^ = 183.5, df = 3, p < 0.001; Table 1). Hosts fed the high-quality S diet had many more spores attacking their gut compared to the poorer quality diets with any amount of *Microcystis*. In general, the Standard clone had more attacking spores in their gut compared to Mid37 (χ^2^ = 19.6, df = 1, p < 0.001). There was no significant interaction between diet and host clone. These results suggest that the variation seen in fungus infection prevalence across different diets (Figure 1a) is at least partially driven by impacts of diet on the earliest stages of infection.

**Figure 2.**
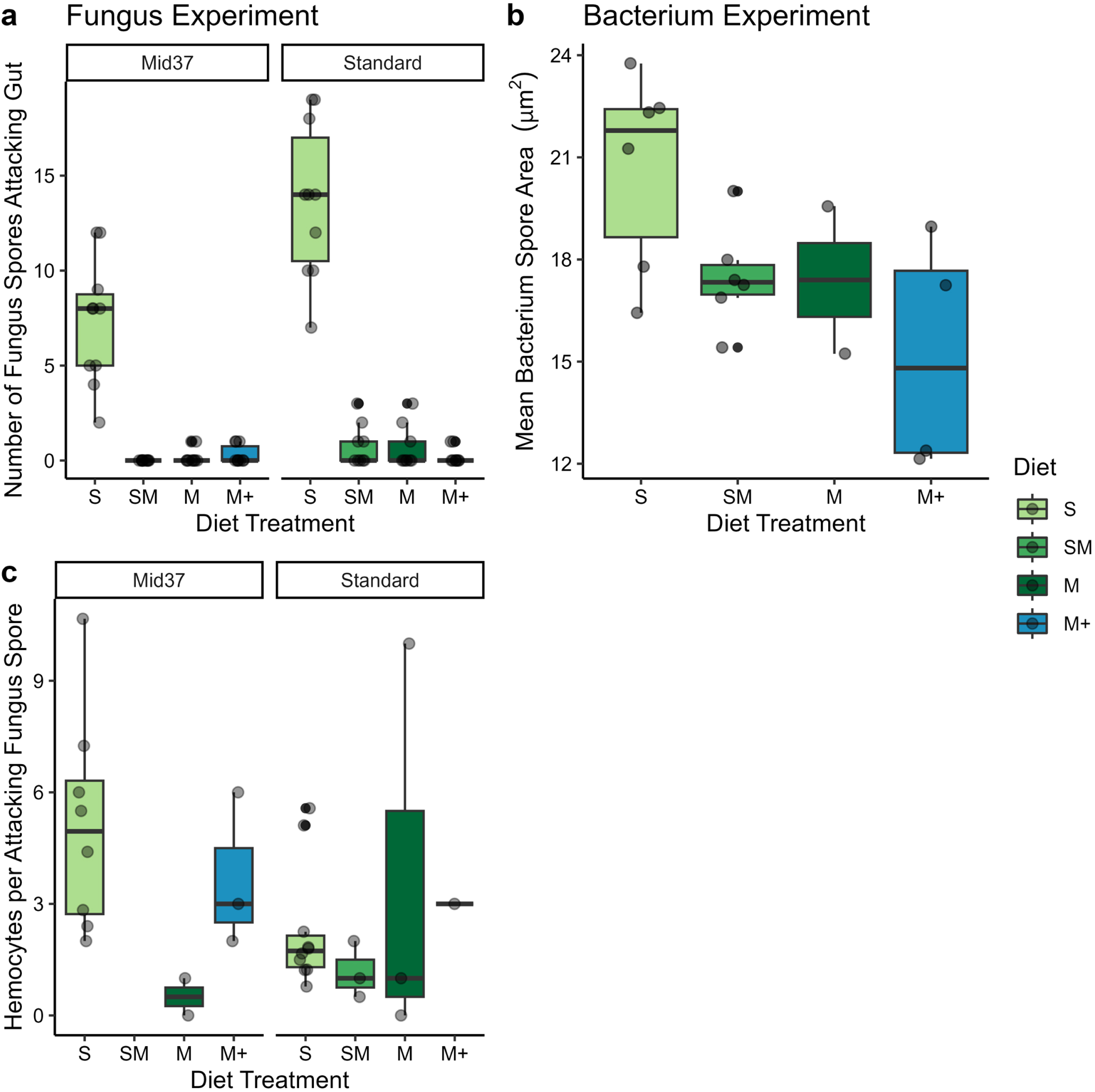
Reducing resource quality reduced the number of attacking spores in the host gut (a), but did not impact host hemocyte immune response (c). Reducing resource quality reduced the size of mature bacterial spores produced (b). a) For both host clones, hosts fed poorer-quality diets that included *Microcystis* (SM, M, and M+) had fewer fungal spores attacking (embedded or penetrating) their gut wall, but Standards had more attacking spores than Mid37s. b) Poorer-quality diets produced smaller mature bacterial spores than high quality S diets. We did not assess spore size differences based on host clone due to Standards relatively lower infection prevalence and spore yields (Figure 1b,f). Bacterial spore size was calculated as the mean area of an ellipse based on two perpendicular diameter measurements of five bacterial spores. c) The number of recruited hemocyte immune cells per fugus spore that punctured the gut wall did not differ based on resource quality diet, but Mid37s recruited more hemocytes per spore than Standards. We were unable to collect data on attacking spores for the bacterium because we do not yet know about the analogous early stages of infection by the bacterium (Ebert et al. 2015). We did not collect data on fungal spore size due to logistic constraints.

Diet did not impact host immune responses, but host clones differed in their investment in immune response (Figure 2c; Table 1). There was no difference in the number of hemocytes per attacking spore based on diet (χ^2^ = 2.6, df = 3, p = 0.45). Mid37s produced more hemocytes per attacking spore compared to Standards (χ^2^ = 4.1, df = 1, p = 0.043).

### Mature parasite spore yield and size

We expected that reduced resource quality might limit the ability of parasites to replicate within hosts. Consistent with this prediction, fungal mature spore yield decreased along with decreasing resource quality (Figure 1e, Table 1, χ^2^ = 19.8, df = 3, p < 0.001); fungal mature spore yield was unaffected by host clone (χ^2^clone = 1.5, df = 1, p = 0.22). In contrast, the bacterial mature spore yield was higher in the Mid37 clone compared to the Standard clone but was unaffected by diet and the diet × clone interaction was not significant (Figure 1f, Table 1, χ^2^diet = 5.6, df = 3, p = 0.14; χ^2^ = 10.91, df = 1, p < 0.001). Consistent with our hypothesis that resource quality may impact the size of each bacterial spore produced, bacterial spore size decreased with reducing resource quality (Figure 2b, F3,14 = 3.6, p = 0.039; Table 1). Log bacterial mature spore yield and average spore size were not significantly correlated (rho = 0.20, p = 0.42).

### Host fecundity and survival: impacts of diet and infection

As expected, diet and host clone strongly influenced host total fecundity (Figure 3a-b, Table 1). In the fungus experiment, both host clones’ fecundity differed based on diet and infection (χ^2^ = 30.1, df = 3, p < 0.0001, χ^2^ = 6.4, df = 2, p = 0.041). Standards had high fecundity that declined sharply with reducing resource quality and infection. Mid37s had similarly low fecundity among the three poorer quality diets (SM, M, and M+) compared to the high-quality S diet; the low overall fecundity and lack of infections in the M and M+ diets make it hard to examine how food quality and infection interacted in this clone (Table S4). In the bacterium experiment, fecundity of the two host clones responded differently to reducing resource quality (χ^2^ = 9.8, df = 3, p = 0.021, Table S6), especially for individuals infected by the bacterium. All bacterium-infected Mid37s produced few offspring regardless of diet.

**Figure 3.**
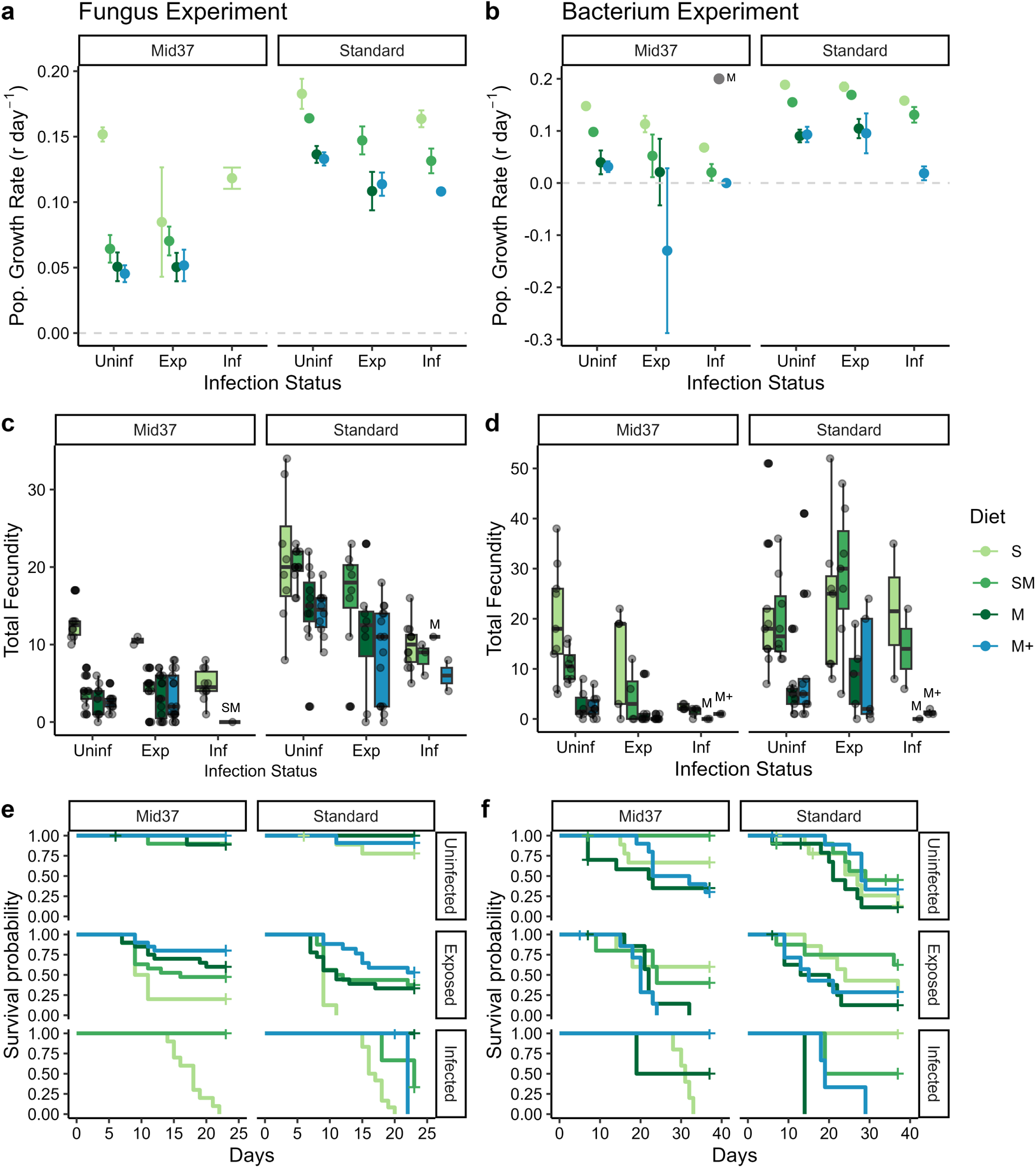
Total fecundity and survival differed based on resource quality diet, host clone, and infection status. a,b) Host lifetime fecundity was strongly impacted by decreasing resource quality diet in the Fungus experiment (a) and Bacterium experiment (b). Total fecundity was calculated as the cumulative sum of all offspring produced during the lifetime of an individual. Note that the y-axis scales are different because the longer bacterium experiment allowed more time for offspring production. c,d) Survival curves for each resource quality diet, host clone, and infection status in the fungus (c) and bacterium (d) experiments. Parasite exposed hosts had lower survival rates compared to uninfected hosts in both experiments. Vertical hashes indicate censored individuals. See the cox proportional hazard ratios in Table S8. Infection status: Uninf = Uninfected (and uninoculated), Exp = Exposed-but-uninfected, Inf = Infected.

However, infected Standards fed higher quality S and SM diets had similar fecundity to uninfected Standards but those fed the poor-quality M and M+ diets had fewer than two offspring on average. Overall, bacterium-infected hosts of both clones produced very few offspring, as would be expected of a castrating parasite, but higher quality diets allowed Standards to still produce offspring.

Host mortality rates differed based on diet and infection status for both parasite experiments and differed between host clones in the fungus experiment (Figure 3c,d, Table S8). In the fungus experiment, infected hosts fed the highest quality S diet died much faster than uninfected and unexposed hosts. In contrast, hosts that were infected with the bacterium had similar mortality rates as uninfected hosts.

### Host fecundity and survival: costs of resisting infection

There was a fecundity cost of resisting fungal infection for Standards but not Mid37s (Table S4; Figure 3a). Fungus-exposed-but-uninfected Standards produced on average 6.6 fewer offspring than uninfected Standards, indicating a fecundity cost of resisting infection; this cost of resisting infection was consistent across diet treatments. In contrast, Mid37s did not show signs of a fecundity cost of resisting fungal infections: uninfected and fungus-exposed-but-uninfected Mid37s had similar fecundity. The two host clones differed in how their fecundity responded to resisting bacterial infection given exposure (χ^2^ = 12.8, df = 2, p = 0.002). Bacterium- exposed-but-uninfected Mid37s produced on average 4.3 fewer offspring than uninfected Mid37s, but there was no difference between exposed-but-uninfected and uninfected Standards (Table S4). For both parasite experiments, the fecundity costs of resisting infection (or lack thereof) were consistent across diets.

Hosts that resisted infection generally paid a high survival cost. Hosts that were fungus-exposed-but-uninfected had a greater than fifteenfold higher risk of death relative to uninfected hosts—but unsurprisingly, infection by the fungus had an even larger (greater than thirty- eightfold) increase in the risk of death (Figure 3c, Table S8). Standards that were fungus-exposed-but-uninfected had a twenty-eightfold increase in risk of mortality compared to uninfected Standards (hazard ratio (HR) = 28.41, p = 0.003), but Mid37s did not have a significantly different risk of death (HR = 6.09, p = 0.11; Table S9). Similarly, bacterium- exposed-but-uninfected hosts had a 67% higher risk of death than uninfected hosts, suggesting that bacterium-exposed hosts also suffer a mortality cost of resisting infection (Figure 3d, Table S8). The Standard’s risk of mortality was not significantly higher than Mid37s, but the two clones did differ based on bacterium exposure. Mid37s that were bacterium-exposed-but-uninfected had 3.25-times (that is, 225% increased) risk of mortality compared to uninfected Mid37s (HR = 3.25, p = 0.0019), while the Standards did not significantly differ in host mortality between uninfected and exposed-but-uninfected hosts (HR = 0.94, p = 0.86; Table S9). For both parasite experiments, the mortality costs of resisting infection were generally consistent across diets. Altogether, the costs of resisting infection varied based on host clone and fungal vs. bacterial inoculation, as summarized in Table S10.

### Body size

As expected, diet strongly influenced host body size in both experiments, with hosts fed the high-quality S diet having the largest body sizes in both experiments (Figure 4). By the end of the experiment, there were signs of parasite-induced gigantism in bacterium-infected Standards, but only in the highest food quality treatment (Figure 4b). This suggests that resource limitation might interfere with the ability of castrating parasites to induce gigantism in their hosts (as also found in Cressler et al. (2014b)). Analyses of host body size during exposure and host growth rate over the experiment are presented in Appendix S1: Section S1.7.

**Figure 4.**
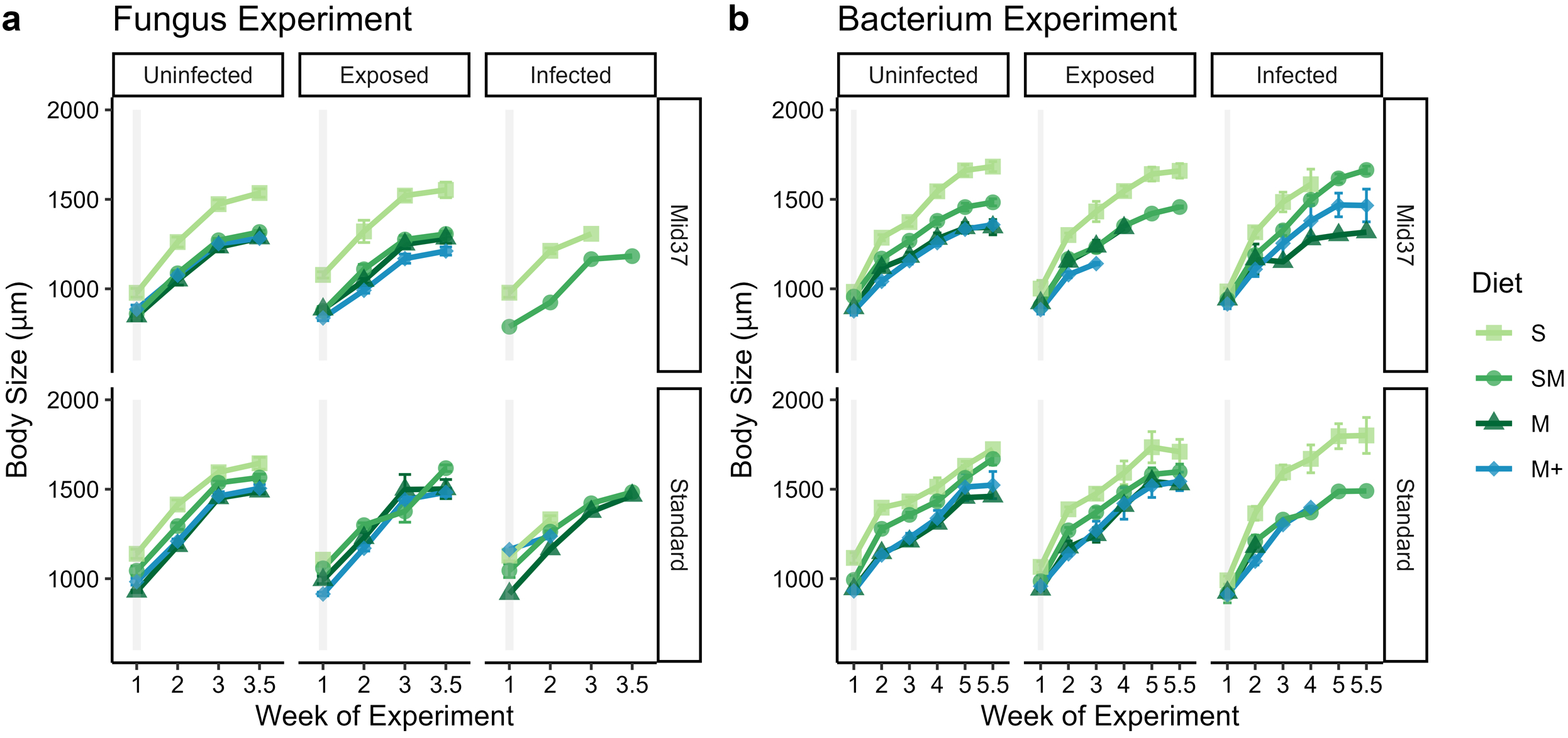
Bacterium-infected Standard individuals showed signs of gigantism, but only in the S treatment, indicating poor quality resources might limit the parasite’s ability to induce gigantism. The y-axis shows the average body size (in micrometers) of individual *Daphnia* over the course of the experiment. Error bars (which are often too small to be visible) represent +/- 1 standard error. Infection status: Uninfected = Uninfected and uninoculated, Exposed = Exposed but uninfected, Infected = Infected. The gray shaded boxes indicate the inoculation period in the first week of the experiment. Some standard error bars are too small to be visible or could not be calculated because there was only a single animal for a given time point and treatment combination.

## Discussion

Resource quality can vary strongly over space and time, both naturally and as a result of human impacts. Here, we sought to mechanistically understand the impact of changing resources on host–parasite interactions, and to see whether those impacts were consistent across host clones and for different parasites. Diet strongly influenced infection prevalence for the fungal parasite but not for the bacterium. These changes in infection prevalence do not seem to be caused by impacts of diet on feeding (and exposure) rate. Instead, for the fungus, the effect of resources on infection prevalence were driven at least in part by impacts of diet on the number of spores that embedded in and penetrated through the host gut wall. Diet did not impact host immune responses (specifically: hemocytes per attacking fungal spore), but immune responses did differ between host clones. Diet influenced spore production, but differently for the two parasites: for the fungus, reduced resource quality limited the number of parasite transmission spores produced, whereas for the bacterium it reduced spore size but not number. Given that the fungal parasite was more sensitive than the bacterium to increasing *Microcystis* in the diet, our results suggest that eutrophication (and the associated increases in *Microcystis* abundance) has the potential to alter the relative abundances of these two parasites.

Diet, host clone, and infection all influenced host fitness. As expected, in the absence of the parasite, ingesting diets with the high quality *Scenedesmus* led to highest fecundity (Figure 5, ‘uninoculated’ orange bars). However, in nature, hosts often face variable resource environments *and* parasites. Because diets with *Microcystis* protect against fungal infections, it was not possible to intuit which diet would maximize fitness in an environment that contains parasites.

**Figure 5.**
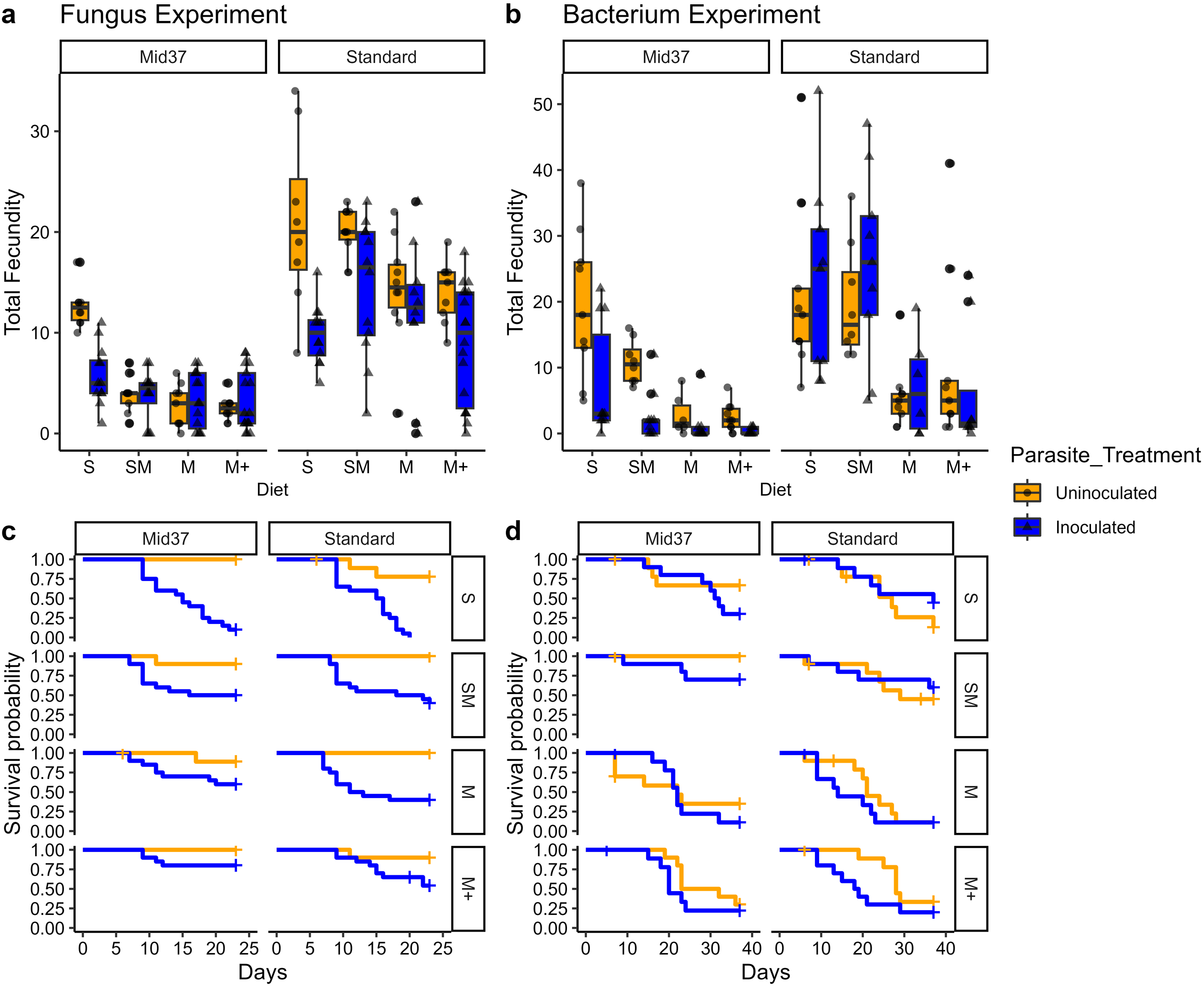
Host reproduction was fairly consistent across diets in the presence of the fungal parasite but not the bacterial parasite. The orange bars show the data for uninoculated and uninfected hosts, and are the same as the ‘Uninf’ bars in Figure 3a&b. The blue bars show the data for all hosts that were inoculated with the fungus (panel a) or bacterium (panel b); thus, these bars combine the data for the ‘Exp’ and ‘Inf’ bars in Figure 3.

Looking at all inoculated hosts (regardless of whether they became infected or resisted infection; Figure 5 ‘inoculated’ blue bars), we can see that the variation in fecundity with diet treatment is greatly reduced in an environment that contains the fungus, but not in one that contains the bacterium. In other words, the protection against the effects of the virulent fungal parasite that diets with *Microcystis* provide were offset by the reduction in fecundity due to the low quality of this diet; because *Microcystis* does not protect against bacterial infections, host fitness remained highest in diets with higher quality *Scenedesmus*.

Reducing resource quality provided hosts protection from the fungal parasite (Figure 1a), but not the bacterial parasite (Figure 1b) despite both parasites sharing several key traits, including environmental transmission through foraging, infection of the hemolymph via exposure through the gut wall, and being obligate killers of the host. This is consistent with an earlier study on this same host species that also found strong effects of diet on infection by the fungus but not the bacterium (Sánchez et al. 2019), though we note that there are other studies that have found effects of diet quality on infection by the bacterium (Frost et al. 2008, Schlotz et al. 2013, Cressler et al. 2014b, Schoebel et al. 2014). In our experiment, hosts had similar filter feeding rates—and therefore, parasite exposure rates—and immune responses across all resource qualities when exposed to the fungus (Figure 1c, Figure 2c), but hosts were smaller and had much lower numbers of attacking fungal spores in their guts when fed diets that included *Microcystis* (SM, M, M+; Figure 2a). *Microcystis* can induce changes in host digestive proteases (von Elert et al. 2012). A recent study found that the digestive protease chymotrypsin promotes dehiscence of fungal spores, which is a key step required for infection (Sánchez et al. 2024). The *Microcystis* CYA43 diet, the same strain used in the present study, strongly suppressed chymotrypsin in a variety of clones of the host, *Daphnia magna* (Sánchez et al. 2024). The Sánchez et al. study proposed that the protective effect of *Microcystis* CYA43 on infection was driven by its suppression of chymotrypsin and the subsequent prevention of spore dehiscence.

Our results in the present study provide further support for this mechanism; it was not possible to quantify attacking spores in the *D. magna* used in the Sánchez et al. study, but here we show that *Microcystis* CYA43 dramatically reduced attacking spores in *D. dentifera*, which is consistent with the diet preventing spore dehiscence. In addition, our data suggest that even diets that are only partially made up of the protective food are sufficient to protect the host from infection. In a study on *D. magna* and a different parasite, white fat cell disease (which is caused by a virus; Toenshoff et al. 2018), there was an approximately linear decrease in infection levels with increasing percentage of *Microcystis* in the diet (Coopman et al. 2014). In our study, the drop off was more dramatic, with a very large reduction in infection levels between 0% and 50% *Microcystis*; there were only modest differences in infection prevalence between 50% and 100% *Microcystis*, though some of this may be because infection levels were nearing the lower bound (of 0% prevalence).

Meanwhile, unlike the fungal parasite, infection by the bacterial parasite was not strongly affected by differences in resource quality in our study. Interestingly, we observed that reducing resource quality tended to reduce parasite exposure via lower feeding rate (Figure 1d), but these differences in exposure rates to the bacterium did not translate into differences in infections. This may be because the bacterial parasite has strong host–parasite genotype specific infectivity, unlike the fungal parasite, where the host and parasite genotypes must match in order for successful infection (Carius et al. 2001, Auld et al. 2012b, Bento et al. 2017, Cornetti et al. 2024). However, in other cases, studies have found an effect of varying food quality on infection by the bacterial parasite, albeit in opposite directions: low quality phosphorus-deficient diets reduced bacterial infections in *D. magna* hosts (Frost et al. 2008), while high quality diets high in polyunsaturated fatty acids protected against infections (Schlotz et al. 2013). Similarly, studies have varied in findings regarding the impact of food quantity on *Pasteuria* infections, with some finding no effect of resource levels on infection (Duneau et al. 2011, Schoebel et al. 2014) and others finding that higher food leads to higher infection (Little et al. 2007, Ben-Ami et al. 2010). An earlier study found that the fecundity reduction caused by *Pasteuria* infection was much larger in hosts fed high quality food, because all hosts (including uninfected ones) had very low fecundity in low resource conditions (Savola and Ebert 2019). Our findings for the Mid37 genotype were consistent with this, but our results for the Standard genotype showed that high food quality allowed it to retain relatively high fecundity even when infected.

We expected that reducing resource quality would reduce the parasite’s capacity to produce new infectious propagules (i.e., spores). We found that the yield of mature fungal spores per individual declined with reducing resource quality (Figure 1e). Prior work has found similar relationships between reduced resource quality and fungal spore yield, where hosts fed poor quality resources were smaller and produced fewer mature spores (Hall et al. 2009, Penczykowski et al. 2014, Manzi et al. 2020) but found conflicting effects on transmission potential. Penczykowski et al. (2014) found that hosts fed poor quality diets were small, had lower exposure rates and produced few mature spores, which all ultimately contributed to lower transmission potential. However, Hall et al. (2009) found the opposite effect, where poor quality diets produce low spore yield, but spores in poor resource quality conditions had a higher probability of infecting new hosts (i.e., transmission). We suggest that these apparently opposing relationships between resource quality and transmission could be due to differences in the identity of the poor-quality phytoplankton, since one study used field seston for the poor-quality treatment while the other used lab reared *Microcystis* (CYA43). Indeed, even just different strains of *Microcystis aeruginosa* can have very different impacts on infection (Tellenbach et al. 2016, Sánchez et al. 2024).

In contrast to our findings for the fungus, the mature spore yield of the bacterial parasite was more strongly affected by host clone than resource quality (Figure 1f). However, there was extreme variation in mature spore yield among individual hosts from greater than 100,000 to less than 1,000, which likely impacted our ability to detect resource quality dependency. Our finding in this study differs from those in a prior study, which found higher bacterial spore yield in hosts fed low quality cyanobacterial diets (Sánchez et al. 2019). In the present study, bacterial spore size did decline with reducing resource quality (Figure 3b), potentially indicating that under poor quality conditions the parasite has limited capacity to energetically invest per spore to maintain adequate spore yields. Other studies have found that reducing food quality or quantity reduces spore yield from infected hosts (Frost et al. 2008, Schlotz et al. 2013, Vale et al. 2013, Nørgaard et al. 2021). However, other studies have not found effects of manipulating resources on bacterial spore yield (Cressler et al. 2014b, Schoebel et al. 2014). These contrasting results highlight that different aspects of resource quality and quantity may produce different outcomes for host–parasite interactions.

We found relatively little difference between the two 100% *Microcystis* diets (one of which was spiked with the microcystin-LR toxin) on host physiology or fitness or infection outcomes in either experiment. At present, the effect of microcystins on susceptibility to *Metschnikowia* is unclear. In one study, diets of *Microcystis* added microcystin-LR had lower infection levels than those with just a non-toxin producing *Microcystis* (Sánchez et al. 2019). However, in a more recent study that used a different strain of *Microcystis* that produces microcystin-LR and that compared it with a mutant that did not, microcystin-LR did not protect against infection (Sánchez et al. 2024). Moreover, two recent studies found that microcystin-LR does not reduce the infectivity of free-living stages of the fungus (Manzi et al. 2022, Sánchez et al. 2023). While the effect of microcystin-LR on infections is not yet resolved, the overall evidence to date also suggests that the primary effect of *Microcystis* on infections is most likely due to its production of protease inhibitors rather than the effect of microcystin-LR (Sánchez et al. 2024). A meta-analysis of laboratory experiments found that the presence of toxins (predominately microcystins) did not negatively affect zooplankton population growth but did reduce zooplankton survival for some toxic *Microcystis* strains (Wilson et al. 2006). In our study, we found that the M and M+ diets had similar fecundity and survival within each host clone and parasite treatment combination. The authors of the meta-analysis study suggest that cyanotoxins like microcystin-LR are not a key driver of poor food quality (Wilson et al. 2006). We propose that the same is likely true of the effect of *Microcystis* on parasitism in *Daphnia*.

Although it would be simpler to assume that all hosts and all parasites will respond to differences in resource quality in the same ways, our study clearly demonstrates that this is not the case. Natural systems are composed of host populations with many host genotypes present, multiple parasites, and fluctuating resource quality and quantity over time and space. Many studies only use a single parasite or single host genotype, but the variation across clones and parasites in our study suggest caution is warranted about making general conclusions. We suggest that the outcomes of host–parasite interactions will likely depend on the specific sensitivities of the host genotypes and parasite species present to resource quality gradients and each host genotype’s degree of susceptibility to different parasites. Therefore, it is essential to conduct studies with multiple genotypes and multiple parasites under realistic environmental conditions to better understand the diversity of host responses that may occur to changing environments and their potential impacts on population dynamics in natural systems.

## Acknowledgements

This work was supported by the US National Science Foundation (DEB-1305836 and DEB-1655856) and the Gordon and Betty Moore Foundation (GBMF9202; https://doi.org/10.37807/GBMF9202). We thank the many members of the Duffy Lab who assisted with conducting the experiments in this study, including Teresa Sauer, Elizabeth Davenport, Kris McIntire, and Marcin Dziuba. We also thank two anonymous reviewers for their helpful feedback on an earlier version of this manuscript.

## Author Contributions

Conceptualization & Study Design: MLF, KFS, S-JS, MAD Data collection: MLF, KFS, S-JS, SKC, KJM, VR Analysis & Visualization: MLF Writing-Original Draft: MLF, MAD Writing-Review & Editing: all authors Resources & Funding: MAD

## References

1. Agrawal, M. K., A. Zitt, D. Bagchi, J. Weckesser, S. N. Bagchi, and E. von Elert. 2005. Characterization of proteases in guts of Daphnia magna and their inhibition by Microcystis aeruginosa PCC 7806. Environmental Toxicology 20:314–322.

2. Auld, S. K., S. R. Hall, J. H. Ochs, M. Sebastian, and M. A. Duffy. 2014. Predators and patterns of within-host growth can mediate both among-host competition and evolution of transmission potential of parasites. American Naturalist 18:S77–S90.

3. Auld, S. K. J. R., A. L. Graham, P. J. Wilson, and T. J. Little. 2012a. Elevated haemocyte number is associated with infection and low fitness potential in wild Daphnia magna. Functional Ecology 26:434–440.

4. Auld, S. K. J. R., S. R. Hall, and M. A. Duffy. 2012b. Epidemiology of a *Daphnia*-multiparasite system and its implications for the Red Queen. PLOS ONE 7:e39564.

5. Auld, S. K. J. R., C. L. Searle, and M. A. Duffy. 2017. Parasite transmission in a natural multihost–multiparasite community. Philosophical Transactions of the Royal Society B: Biological Sciences 372.

6. Bates, D., M. Maechler, B. Bolker, and S. Walker. 2015. Fitting linear mixed-effects models using lme4. Journal of Statistical Software 67:1–48.

7. Becker, D. J., and R. J. Hall. 2014. Too much of a good thing: resource provisioning alters infectious disease dynamics in wildlife. Biology Letters 10:20140309.

8. Beldomenico, P. M., S. Telfer, S. Gebert, L. Lukomski, M. Bennett, and M. Begon. 2008. Poor condition and infection: a vicious circle in natural populations. Proceedings of the Royal Society B: Biological Sciences 275:1753–1759.

9. Ben-Ami, F., D. Ebert, and R. R. Regoes. 2010. Pathogen Dose Infectivity Curves as a Method to Analyze the Distribution of Host Susceptibility: A Quantitative Assessment of Maternal Effects after Food Stress and Pathogen Exposure. American Naturalist 175:106–115.

10. Bento, G., J. Routtu, P. D. Fields, Y. Bourgeois, L. Du Pasquier, and D. Ebert. 2017. The genetic basis of resistance and matching-allele interactions of a host-parasite system: The *Daphnia magna-Pasteuria ramosa* model. PLOS Genetics 13:e1006596.

11. Brooks, M. E., K. Kristensen, K. J. V. Benthem, A. Magnusson, C. W. Berg, A. Nielsen, H. J. Skaug, M. Mächler, and B. M. Bolker. 2017. glmmTMB Balances Speed and Flexibility Among Packages for Zero-inflated Generalized Linear Mixed Modeling. The R Journal 9:378–400.

12. Budischak, S. A., and C. E. Cressler. 2018. Fueling Defense: Effects of Resources on the Ecology and Evolution of Tolerance to Parasite Infection. Frontiers in Immunology 9.

13. Burberg, C., T. Petzoldt, and E. von Elert. 2020. Phosphate Limitation Increases Content of Protease Inhibitors in the Cyanobacterium Microcystis aeruginosa. Toxins 12:33.

14. Cáceres, C. E., and T. E. Stewart Merrill. 2023. The role of varying resources on *Daphnia dentifera* immune responses. Fundamental and Applied Limnology 196:217–228.

15. Carius, H. J., T. J. Little, and D. Ebert. 2001. Genetic variation in a host-parasite association: Potential for coevolution and frequency-dependent selection. Evolution 55:1136–1145.

16. Carpenter, S. R. 2008. Phosphorus control is critical to mitigating eutrophication. Proceedings of the National Academy of Sciences 105:11039–11040.

17. Chaffin, J. D., J. A. Westrick, L. A. Reitz, and T. B. Bridgeman. 2023. Microcystin congeners in Lake Erie follow the seasonal pattern of nitrogen availability. Harmful Algae 127:102466.

18. Chorus, I., and M. Welker, editors. 2021. Toxic Cyanobacteria in Water: A guide to their public health consequences, monitoring and management. CRC Press, London.

19. Civitello, D. J., B. E. Allman, C. Morozumi, and J. R. Rohr. 2018. Assessing the direct and indirect effects of food provisioning and nutrient enrichment on wildlife infectious disease dynamics. Philosophical Transactions of the Royal Society B: Biological Sciences 373:20170101.

20. Coopman, M., K. Muylaert, B. Lange, L. Reyserhove, and E. Decaestecker. 2014. Context dependency of infectious disease: the cyanobacterium Microcystis aeruginosa decreases white bacterial disease in Daphnia magna. Freshwater Biology 59:714–723.

21. Cornet, S., C. Bichet, S. Larcombe, B. Faivre, and G. Sorci. 2013. Impact of host nutritional status on infection dynamics and parasite virulence in a bird-malaria system. Journal of Animal Ecology:n/a-n/a.

22. Cornetti, L., P. D. Fields, L. Du Pasquier, and D. Ebert. 2024. Long-term balancing selection for pathogen resistance maintains trans-species polymorphisms in a planktonic crustacean. Nature Communications 15:5333.

23. Cotter, S. C., and E. Al Shareefi. 2022. Nutritional ecology, infection and immune defence — exploring the mechanisms. Current Opinion in Insect Science 50:100862.

24. Cressler, C. E., W. A. Nelson, T. Day, and E. McCauley. 2014a. Disentangling the interaction among host resources, the immune system and pathogens. Ecology Letters 17:284–293.

25. Cressler, C. E., W. A. Nelson, T. Day, and E. McCauley. 2014b. Starvation reveals the cause of infection-induced castration and gigantism. Proceedings of the Royal Society B: Biological Sciences 281.

26. Duffy, M. A., and L. Sivars-Becker. 2007. Rapid evolution and ecological host-parasite dynamics. Ecology Letters 10:44–53.

27. Duneau, D., P. Luijckx, F. Ben-Ami, C. Laforsch, and D. Ebert. 2011. Resolving the infection process reveals striking differences in the contribution of environment, genetics and phylogeny to host-parasite interactions. BMC Biology 9:11.

28. Ebert, D. 2005. Ecology, epidemiology and evolution of parasitism in *Daphnia*. National Library of Medicine (US), National Center for Biotechnology Information, Bethesda, MD.

29. Ebert, D., D. Duneau, M. D. Hall, P. Luijckx, J. P. Andras, L. Du Pasquier, and F. Ben-Ami. 2015. A population biology perspective on the stepwise infection process of the bacterial pathogen *Pasteuria ramosa* in *Daphnia*. Advances in Parasitology in press.

30. Ebert, D., M. Lipsitch, and K. L. Mangin. 2000. The effect of parasites on host population density and extinction: Experimental epidemiology with *Daphnia* and six microparasites. American Naturalist 156:459–477.

31. Fox, J., and S. Weisberg. 2019. An {R} Companion to Applied Regression, Third Edition. Sage, Thousand Oaks, CA.

32. Frost, P. C., D. Ebert, and V. H. Smith. 2008. Responses of a bacterial pathogen to phosphorus limitation of its aquatic invertebrate host. Ecology 89:313–318.

33. Grobbelaar, J. U. 2004.Algal nutrition: mineral nutrition.

34. Ha, J. H., T. Hidaka, and H. Tsuno. 2009. Quantification of Toxic Microcystis and Evaluation of Its Dominance Ratio in Blooms Using Real-Time PCR. Environmental Science & Technology 43:812–818.

35. Hall, S. R., C. R. Becker, M. A. Duffy, and C. E. Cáceres. 2012. A power-efficiency tradeoff in resource use alters epidemiological relationships. Ecology 93:645–656.

36. Hall, S. R., C. Knight, C. R. Becker, M. A. Duffy, A. J. Tessier, and C. E. Cáceres. 2009. Quality matters: resource quality for hosts and the timing of epidemics. Ecology Letters 12:118–128.

37. Hartig, F. 2020. DHARMa: Residual diagnostics for hierarchical (multi-level/mixed) regression models. https://cran.r-project.org/package=DHARMa.

38. Hayes, N. M., H. A. Haig, G. L. Simpson, and P. R. Leavitt. 2020. Effects of lake warming on the seasonal risk of toxic cyanobacteria exposure. Limnology and Oceanography Letters 5:393–402.

39. Heuermann, N., F. van Langevelde, S. E. van Wieren, and H. H. T. Prins. 2011. Increased searching and handling effort in tall swards lead to a Type IV functional response in small grazing herbivores. Oecologia 166:659–669.

40. Hite, J. L., A. C. Pfenning-Butterworth, R. E. Vetter, and C. E. Cressler. 2020. A high- throughput method to quantify feeding rates in aquatic organisms: A case study with Daphnia. Ecol Evol 10:6239–6245.

41. Huang, Y., P. Li, G. Chen, L. Peng, and X. Chen. 2018. The production of cyanobacterial carbon under nitrogen-limited cultivation and its potential for nitrate removal. Chemosphere 190:1–8.

42. Huffman, M. A., and J. M. Caton. 2001. Self-induced Increase of Gut Motility and the Control of Parasitic Infections in Wild Chimpanzees. International Journal of Primatology 22:329–346.

43. Huisman, J., G. A. Codd, H. W. Paerl, B. W. Ibelings, J. M. H. Verspagen, and P. M. Visser. 2018. Cyanobacterial blooms. Nature Reviews Microbiology 16:471–483.

44. Iannino, A., P. Fink, A. T. L. Vosshage, and M. Weitere. 2023. Resource-dependent foraging behaviour of grazers enhances effects of nutrient enrichment on algal biomass. Oecologia 201:479–488.

45. Ibelings, B. W., K. Bruning, J. de Jonge, K. Wolfstein, L. M. Pires, J. Postma, and T. Burger. 2005. Distribution of microcystins in a lake foodweb: no evidence for biomagnification. Microb Ecol 49:487–500.

46. Kassambara, A., M. Kosinski, and P. Biecek. 2011. survminer: Drawing Survival Curves using ’ggplot2’. R package version 0.4.9.

47. Klüttgen, B., U. Dulmer, M. Engels, and H. T. Ratte. 1994. ADaM, an artificial freshwater for the culture of zooplankton. Water Research 28:743–746.

48. Kutzer, M. A. M., V. Gupta, K. Neophytou, V. Doublet, K. M. Monteith, and P. F. Vale. 2023. Intraspecific genetic variation in host vigour, viral load and disease tolerance during Drosophila C virus infection. Open Biology 13:230025.

49. Leitão, A. B., X. Bian, J. P. Day, S. Pitton, E. Demir, and F. M. Jiggins. 2019. Independent effects on cellular and humoral immune responses underlie genotype-by-genotype interactions between Drosophila and parasitoids. PLOS Pathogens 15:e1008084.

50. Lenth, R. V. 2022. emmeans: Estimated Marginal Means, aka Least-Squares Means. R package version 1.7.3.

51. Little, T., J. Birch, P. Vale, and M. Tseng. 2007. Parasite transgenerational effects on infection. Evolutionary Ecology Research 9:459–469.

52. Luijckx, P., F. Ben-Ami, L. Mouton, L. Du Pasquier, and D. Ebert. 2011. Cloning of the unculturable parasite Pasteuria ramosa and its Daphnia host reveals extreme genotype– genotype interactions. Ecology Letters 14:125–131.

53. Manzi, F., R. Agha, Y. Lu, F. Ben-Ami, and J. Wolinska. 2020. Temperature and host diet jointly influence the outcome of infection in a Daphnia-fungal parasite system. Freshwater Biology 65:757-767.

54. Manzi, F., R. Agha, M. Mühlenhaupt, and J. Wolinska. 2022. Prior exposure of a fungal parasite to cyanobacterial extracts does not impair infection of its Daphnia host. Hydrobiologia.

55. McLean, K. D. 2022. Ecological and evolutionary drivers of host defenses and pathogen infectivity shape host-parasite interactions at multiple levels of biological organization. University of Michigan, Ann Arbor.

56. Mella, V. S. A., M. Possell, S. M. Troxell-Smith, and C. McArthur. 2018. Visit, consume and quit: Patch quality affects the three stages of foraging. Journal of Animal Ecology 87:1615–1626.

57. Mugabo, M., D. Gilljam, L. Petteway, C. Yuan, M. S. Fowler, and S. M. Sait. 2019. Environmental degradation amplifies species’ responses to temperature variation in a trophic interaction. Journal of Animal Ecology 88:1657–1669.

58. Nørgaard, L. S., G. Ghedini, B. L. Phillips, and M. D. Hall. 2021. Energetic scaling across different host densities and its consequences for pathogen proliferation. Functional Ecology 35:475–484.

59. Oro, D., M. Genovart, G. Tavecchia, M. S. Fowler, and A. Martínez-Abraín. 2013. Ecological and evolutionary implications of food subsidies from humans. Ecology Letters 16:1501–1514.

60. Penczykowski, R. M., B. C. P. Lemanski, R. D. Sieg, S. R. Hall, J. Housley Ochs, J. Kubanek, and M. A. Duffy. 2014. Poor resource quality lowers transmission potential by changing foraging behaviour. Functional Ecology 28:1245–1255.

61. R Core Team. 2022. R: A Language and Environment for Statistical Computing. R Foundation for Statistical Computing, Vienna, Austria.

62. Robb, G. N., R. A. McDonald, D. E. Chamberlain, and S. Bearhop. 2008. Food for thought: supplementary feeding as a driver of ecological change in avian populations. Frontiers in Ecology and the Environment 6:476–484.

63. Sánchez, K. F., N. Huntley, M. A. Duffy, and M. D. Hunter. 2019. Toxins or medicines? Phytoplankton diets mediate host and parasite fitness in a freshwater system. Proceedings of the Royal Society B 286:20182231.

64. Sánchez, K. F., E. von Elert, K. Monell, S. K. Calhoun, A. Maisha, P. McCreadie, and M. A. Duffy. 2024. Inhibition of gut digestive proteases by cyanobacterial diets decreases infection in a Daphnia host-parasite system. Authorea.

65. Sánchez, K. F., B. Zhong, J. A. Agudelo, and M. A. Duffy. 2023. Infectivity of the parasite Metschnikowia bicuspidata is decreased by time spent as a transmission spore, but exposure to phycotoxins in the water column has no effect. Freshwater Biology 68:1020–1030.

66. Sandland, G. J., and D. J. Minchella. 2003. Costs of immune defense: an enigma wrapped in an environmental cloak? Trends in Parasitology 19:571–574.

67. Savola, E., and D. Ebert. 2019. Assessment of parasite virulence in a natural population of a planktonic crustacean. BMC Ecology 19:14.

68. Schatz, G. S., and E. McCauley. 2007. Foraging behavior by Daphnia in stoichiometric gradients of food quality. Oecologia 153:1021–1030.

69. Schlotz, N., D. Ebert, and D. Martin-Creuzburg. 2013. Dietary supply with polyunsaturated fatty acids and resulting maternal effects influence host--parasite interactions. BMC Ecol 13:41.

70. Schoebel, C. N., S. Auld, P. Spaak, and T. J. Little. 2014. Effects of Juvenile Host Density and Food Availability on Adult Immune Response, Parasite Resistance and Virulence in a Daphnia-Parasite System. PLOS ONE 9:9.

71. Schwarzenberger, A., C. J. Kuster, and E. Von Elert. 2012. Molecular mechanisms of tolerance to cyanobacterial protease inhibitors revealed by clonal differences in Daphnia magna. Molecular Ecology:n/a-n/a.

72. Searle, C. L., J. H. Ochs, C. E. Cáceres, S. L. Chiang, N. M. Gerardo, S. R. Hall, and M. A. Duffy. 2015. Plasticity, not genetic variation, drives infection success of a fungal parasite. Parasitology 142:839–848.

73. Smucker, N. J., J. J. Beaulieu, C. T. Nietch, and J. L. Young. 2021. Increasingly severe cyanobacterial blooms and deep water hypoxia coincide with warming water temperatures in reservoirs. Global Change Biology 27:2507–2519.

74. Stevenson, M., T. Nunes, C. Heuer, J. Marshall, J. Sanchez, R. Thornton, J. Reiczigel, J. Robison-Cox, P. Sebastiani, P. Solymos, K. Yoshida, G. Jones, S. Pirikahu, S. Firestone, R. Kyle, J. Popp, M. Jay, and C. Reynard. 2020. epiR:Tools for the analysis of epidemiological data.

75. Stewart Merrill, T. E., and C. E. Cáceres. 2018. Within-host complexity of a plankton-parasite interaction. Ecology 99:2864–2867.

76. Stewart Merrill, T. E., S. Hall, L. Merrill, and C. E. Cáceres. 2019. Variation in immune defense shapes disease outcomes in laboratory and wild Daphnia. Integrative and Comparative Biology.

77. Sun, S.-J., M. K. Dziuba, R. N. Jaye, and M. A. Duffy. 2023. Temperature modifies trait- mediated infection outcomes in a *Daphnia*-fungal parasite system. Philosophical Transactions of the Royal Society B: Biological Sciences 378:20220009.

78. Tellenbach, C., N. Tardent, F. Pomati, B. Keller, N. G. Hairston, J. Wolinska, and P. Spaak. 2016. Cyanobacteria facilitate parasite epidemics in Daphnia. Ecology 97:3422–3432.

79. Tessier, A. J., and P. Woodruff. 2002. Cryptic trophic cascade along a gradient of lake size. Ecology 83:1263–1270.

80. Therneau, T. 2022. A Package for Survival Analysis in R_. R package version 3.3-1.

81. Therneau, T., and P. M. Grambsch. 2000. Modeling survival data: Extending the Cox model. Springer, New York.

82. Toenshoff, E. R., P. D. Fields, Y. X. Bourgeois, and D. Ebert. 2018. The End of a 60-Year Riddle: Identification and Genomic Characterization of an Iridovirus, the Causative Agent of White Fat Cell Disease in Zooplankton. G3: Genes|Genomes|Genetics.

83. Vale, P. F., M. Choisy, and T. J. Little. 2013. Host nutrition alters the variance in parasite transmission potential. Biology Letters 9:20121145.

84. Watt, W. B. 1986. Power and Efficiency as Indexes of Fitness in Metabolic Organization. The American Naturalist 127:629–653.

85. Weterings, M. J. A., S. Moonen, H. H. T. Prins, S. E. van Wieren, and F. van Langevelde. 2018. Food quality and quantity are more important in explaining foraging of an intermediate-sized mammalian herbivore than predation risk or competition. Ecology and Evolution 8:8419–8432.

