## Appendix for "Resource Quality Differentially Impacts *Daphnia* Interactions with Two Parasites"

### Appendix S1

### Section S1 Additional Methods and Results

#### *S1.1 Analysis of microcystin-LR concentrations among treatments and between experiments*

We evaluated whether there were any differences in microcystin-LR concentration among the four diets and between the two experiments. We used a linear regression on the log + 1 transformed microcystin concentration collected during parasite exposure on day 8 for all treatments in both experiments. We tested resource quality diet and experiment as predictors, and the interaction between these two factors. We conducted two follow up post-hoc tests to evaluate the specific differences among levels: 1) differences between each experiment conditional on diet treatment, and 2) differences among diets conditional on the experiment. We found no differences in microcystin concentration based on experiment ($F_{1,96} = 1.998$, $p = 0.16$), and there was no significant interaction between experiment and resource quality diet ($F_{3,96} = 0.41$, $p = 0.74$). We also found no difference between the two experiments when examining within each diet treatment. Unsurprisingly, we did find a strongly significant effect of resource quality diet (Figure S1; $F_{3,96} = 1444.06$, $p < 0.0001$), since the diet treatments determined which animals received the M+ toxin spiked diet. Importantly, the M+ toxin spiked treatment had significantly higher microcystin-LR concentrations by about an order of magnitude over all the other diets (S, SM, and M). There were no differences between the three S, SM, and M diets that did not have microcystin added, indicating that neither the *Scenedesmus* nor *Microcystis* phytoplankton contributed to microcystin concentration in the beakers. The low level of microcystin concentration found across all S, SM, and M beakers therefore most likely represents the concentration found in the filtered lake water used throughout the experiment.

There was an accidental 'super spike' of microcystin on day 4 of the bacterium experiment, as described in the main text; animals were moved out of this super-spiked water on day 5. We compared the microcystin-LR concentrations of the M and M+ diets at the end of the toxin spike at day 5 (just prior to when animals were moved to water with the intended amount of microcystin-LR) and during parasite exposure at day 8 in the bacterium experiment to assess the relative differences in microcystin concentration. We used a linear regression on the log of microcystin concentration with experiment day (day 5 or 8) and resource quality diet (M or M+), and their interaction as the main predictors. Two outliers were removed from day 5 to improve the normality of the residuals. Both main factors and the day×diet interaction were significant (Figure S2; $F_{1,32} = 27.83$, $p < 0.0001$). A post-hoc test using emmeans allowed us to investigate the interaction, which showed that the microcystin-LR concentration in M+ during the toxin spike on day 5 was significantly higher than that of M+ on day 8 during parasite exposure ($p < 0.0001$), but there was no difference between in the microcystin concentrations of the two M diets on day 5 and day 8 ($p = 0.91$).

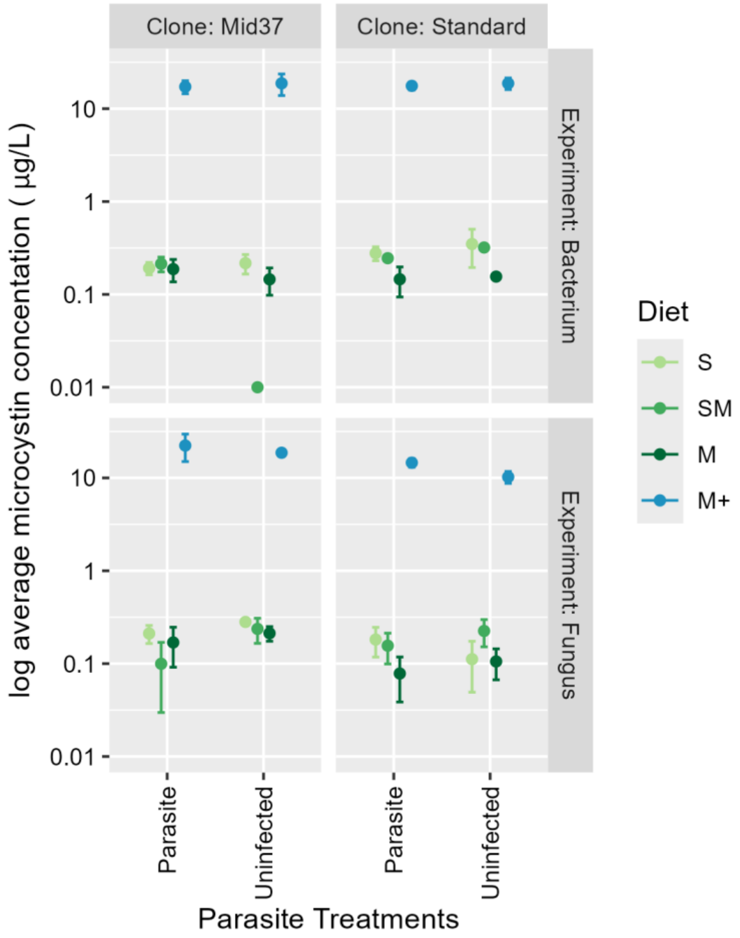


**Figure S1.** Microcystin-LR concentrations (µg/L) during parasite exposure on Day 8 of the fungus and bacterium experiments for each resource quality diet × parasite × clone treatment. The 100% *Microcystis* with spiked microcystin (M+) diet had consistently higher microcystin-LR concentrations than all other diets. There was no difference in microcystin-LR concentrations among the S, SM, or M diets within an experiment ($p_{bacterium} > 0.71$; $p_{fungus} > 0.91$) but M+ was significantly higher than all other diets ($p < 0.001$). There was no difference between experiments for each paired diet treatment ($F_{1,96} = 1.998$, $p = 0.16$).

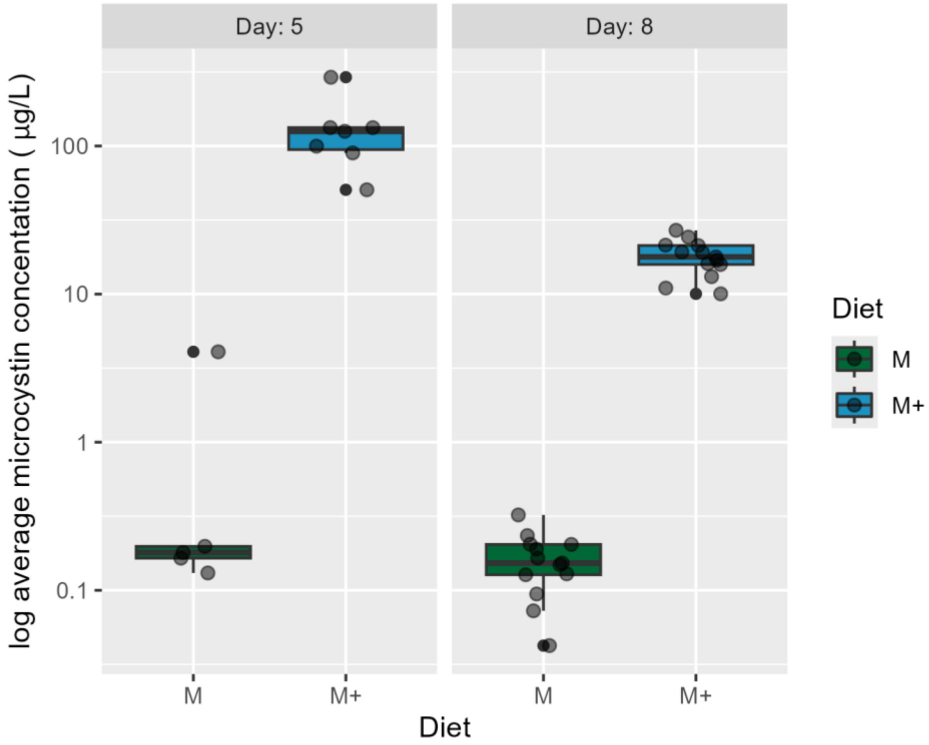


**Figure S2.** For the bacterium experiment, we accidentally added extra microcystin-LR to the M+ spiked toxin treatment ("super spike") on day 4, which was before parasite exposure. Animals were in this water for one day. We show the difference in microcystin-LR concentrations (µg/L) during the accidental super spike (water collected on Day 5 just before animals were moved to the water with the intended amount of microcystin-LR) and during parasite exposure (Day 8) for the 100% *Microcystis* (M) and 100% *Microcystis* + toxin (M+) treatments. There was a significant interaction between diet and experimental day 5 and 8 ($F_{1,32} = 27.83$, $p < 0.0001$), where the microcystin concentration in the M+ diet on day 5 was significantly higher than M+ diet on day 8 ($p < 0.001$), but there was no difference between the two M diets on day 5 and day 8 ($p = 0.91$).

*S1.2 Additional methods for offspring production, infection status, and spore yield*

During each water change we recorded the number of offspring produced and infection status. For each host, we summed offspring produced each week of the experiment and the total number of offspring produced during the experiment ('total fecundity'). Hosts were visually assessed for symptomatic infections under a dissecting scope, which were later confirmed with spore counts. Hosts that produced mature spores were considered 'infected', which is equivalent to the 'terminal infection' designation of Stewart Merrill et al. (2019). Hosts that died during the experiment were saved individually in 1.5 mL microcentrifuge tubes with 0.1 mL deionized water and stored at -20°C for spore counts. To determine spore yield per host, we ground up individual hosts with an electric pestle for 60 s to homogenize the spore solution. We pipetted 10 µL of the spore solution onto a Neubauer Hemocytometer and averaged the counts of mature spores from four grids and multiplied by 10,000 (per manufacturer's instructions to get count per mL) and 0.1-mL volume to estimate the number of mature spores per host. Inoculated hosts that died within 7 days from the start of parasite exposure (i.e., experiment day 14 and prior) were excluded from infection prevalence and mature spore yield analyses (excluded: $n_{fungus}$ = 64, $n_{bacterium}$ = 25). Hosts that did not produce any mature spores were excluded from the mature spore yield analyses; only one host in each experiment contained immature spores but not mature spores.

*S1.3 Additional methods for spore size*

We also measured spore size for the bacterial parasite while counting spores on the hemocytometer by measuring the spore diameter in two perpendicular directions using cellSens Standard Software (Olympus, version 1.18). Five spores were measured per infected host, then we calculated the surface area of an oval and averaged the spore surface area per individual host. We did not measure spore size for animals with estimated mature spore count of less than 250 for the entire animal because we could not get sufficient replicate measurements on the hemocytometer. Due to logistic constraints, we did not take spore size measurements for the fungus.

*S1.4 Additional feeding rate and growth rate methods*

We measured host feeding rate, respiration rate, and body size during parasite exposure and each week of the experiment thereafter (bacterium experiment: five measurements total; fungus experiment: three measurements total). Due to the number of replicates and the length of these protocols, we ran the feeding and respiration rate trials over two days. Each treatment combination (N = 10) was split into two equal-size blocks of 5 replicates (e.g., Block 1: S Mid37 Uninfected Reps 1 – 5, Block 2: S Mid37 Uninfected Reps 6 – 10). On the first day of parasite exposure, Block 1 completed the feeding rate assay and body length measurements, while Block 2 completed the respiration rate assay; the following day, each block completed the other assay.

We used the feeding rate assay to assess individual host's energetic intake under different resource quality conditions over time and parasite exposure. Prior work has shown that *D. dentifera* feeding rates are strongly correlated with consumption of parasite spores while *D. dentifera* are filter feeding in the water column (Shocket et al. 2018, Strauss et al. 2019). Additionally, *D. dentifera* alter their filtering rate based on their environmental conditions, including the abundance and type of phytoplankton in the water column (Penczykowski et al. 2014). We followed the feeding rate protocol described in Hite et al. (2020). In brief, for each resource quality treatment we created a homogenized 300 mL stock beaker of filtered lake water with 2 mg/L of the appropriate diet food and 15 ug/L Microcystin-LR for the M+ treatment added (as described above). We filled glass 15 mL falcon tubes with 10 mL of the homogenized water from each resource quality stock beaker and placed individual hosts according to their diet treatment into the falcon tubes and capped tightly. For each diet treatment, we also included 'no *Daphnia*' and 'no algae' control tubes. The 'no *Daphnia*' control received the same 2 mg/L concentration of algae as the rest of the diet treatment but did not have a host added to the tube. The 'no algae' control only included 10 mL of filtered lake water without any phytoplankton added. We then allowed the hosts to feed for 6 h, while gently inverting each tube every 30 min to resuspend the algae. The entire assay (set up to take down) was conducted in the dark. After 6 h of feeding, we took eight 200 µL subsamples of water from each tube and placed them in a black 96-well plate (Microfluor 1, Thermo Fisher Scientific, Waltham, MA). Each plate included eight subsamples of the 'no *Daphnia*' and 'no algae' controls each. We measured endpoint fluorescence intensity of chlorophyll *a* using top read mode at 435 nm excitation and 665 nm emission wavelengths during a 10-second orbital shake at 22° C in a plate reader (Biotek Synergy H1, Aligent, Santa Clara, CA). All water subsamples were averaged together, with individual well readings that were statistical outliers compared to other readings within the same diet treatment excluded from the average (N = 10; Rosner Test, *EnvStats* package (Millard 2013)).

We measured the body size of each animal after the feeding rate assay to be able to account for differences in feeding behavior based on relative body size, since larger *Daphnia* have higher filtering rates following a square function with body length (Hall et al. 2007). Since animals fed higher quality diets were expected to grow faster than animals fed poor quality diets, we analyzed both standard feeding behavior and body size-controlled feeding behavior to evaluate whether body size impacted how hosts responded to the diet and parasite treatments. We measured the length of each animal under a dissecting microscope with 4× magnification by drawing a straight line from the middle of the *Daphnia*'s eye to the base of their tail spine on the ventral side. All measurements were taken with cellSens Standard Software (Olympus, version 1.18). We calculated the body size growth rate per day using the difference in the body length measurements from the parasite exposure (Day 7 and 8) and the following week (Day 14 and 15) divided by seven days.

We calculated the feeding rate of individual *D. dentifera*, $f$ (mL ind$^{-1}$ h$^{-1}$), based on derived formulas described in Hite et al. (2020), as the difference in fluorescence between *D. dentifera* experimental tubes and 'no *Daphnia*' control tubes from the same plate:

$$f = \ln \frac{F_{control}}{F_{Daphnia}} \times \frac{V}{t},$$

where $F_{control}$ is average fluorescence of control wells, $F_{Daphnia}$ is the average fluorescence of treatment wells that contained an animal, $V$ is the volume of the filtered lake water and appropriate diet food in the feeding tube (10 mL), and $t$ is the time that the *Daphnia* fed in hours (approximately 6 h). We excluded animals that died during the feeding rate assay, or within 24 h of the feeding rate assay conducted during parasite exposure (excluded: $n_{fungus}$ = 11 (6.9%), $n_{bacterium}$ = 10 (6.25%); feeding rate was not measured on the additional replicates for the fungus experiment). We also calculated the body size corrected feeding rate (relative feeding rate) of individual *D. dentifera* (mL ind$^{-1}$ h$^{-1}$ mm$^{-2}$) by dividing the calculated feeding rate by the square of the length of the *Daphnia* in millimeters.

#### *S1.4 Additional metabolic rate methods*

We measured the metabolic rate during each week of the experiment following the schedule for the two blocks described above to estimate individual host's energetic expenditure under different resource quality conditions over time and parasite exposure. Specifically, we measured oxygen consumption rates ($VO_2$) as a proxy for metabolic rates (White 2011) using a fluorescence-based respirometry system following Nørgaard et al. (2021). Energy expenditure has been shown to change with host ingestion rates, activity, and stress levels (e.g., Bergman Filho et al. 2011, Hall et al. 2024), and the difference between energy intake and expenditure will alter the discretionary energy available for hosts and parasites. We placed individual animals from each treatment into 2 mL glass vials with oxygen sensor spots on the bottom and filled with room temperature, autoclaved filtered lake water with 2 mg/L of the appropriate diet food and 15 µg/L microcystin-LR for the M+ treatment added (as described above). For each diet treatment, we also included four blank control vials with the same concentration of the appropriate diet food, but no animal was added to the vials. For the bacterium and fungus experiments, all experimental animal vials and the four blank control vials for a given diet treatment were grouped together on a single 24-well plate. For each respiration day, we conducted two 2-h and 20-min runs of the respiration assay using two 24-channel PreSens oxygen readers (SDR SensorDish Reader, Precision Sensing GmbH, Germany) such that all four diet treatment plates for a block were run with two diet treatments run simultaneously. Each assay measured percent air saturation every two minutes and maintained in a temperature-controlled room at 22° C in the dark. Experimental animals were placed back in their original beakers after the respiration assay. The entire respiration assay set up and take down were completed in the dark, and vials were cleaned with 10% ethanol solution between assays. We calculated the rate of oxygen consumption (VO2, mL O2/hr) using the change in oxygen saturation over time (%/hr) following prior studies (Alton et al. 2007, Nørgaard et al. 2021):

$$VO_2 = -1 \times \left[\left(m_{Daphnia} - \frac{m_{control}}{100}\right)\right] \times V \times \beta O_2,$$

where $m_{Daphnia}$ is the rate of change in the oxygen saturation of experimental animal vials, $m_{control}$ is the per run average rate of change for the blank controls, $V$ is the volume of filtered lake water in the vials (0.002 L), and $\beta O_2$ is the oxygen capacitance of air-saturated water at 22° C (6.15 mL $O_2$ per L of water (Cameron 1986)). We estimated the monotonic slope for each experimental animal and control vial to get $m_{Daphnia}$ and $m_{control}$ parameters, respectively, using the LoLinR package in R (Olito et al. 2017). We then calculated the metabolic rate (J/hr) from the VO2 estimates (mL O2/hr) using the calorific conversion factor of 20.13 J/mL O2 (Lighton 2008). We excluded metabolic rates for animals that died during the respiration rate assay and those that died within 24 h of the respiration trial that was conducted during parasite exposure (excluded: $n_{fungus}$ = 10 (6.25%), $n_{bacterium}$ = 15 (9.4%); respiration rate was not measured on the additional replicates for the fungus experiment). One sample with a bad VO2 reading (S Uninfected Mid37 Rep 9) and one pair of mislabeled samples that could not be distinguished (M Uninfected Mid37 and Standard Rep 7s) were removed from the metabolic rate analyses. Additionally, 3 blank vials with extreme monotonic slopes were excluded from the calculations ($N_{bacterium}$ = 1, $N_{fungus}$ = 2).

*S1.5 Results for mature spore yield and microcystin-LR concentration*

We ran GLMMs for mature fungal and bacterial spore yield with a quasipoisson distribution and a log link function to test the effect of average microcystin-LR concentration on spore yield while controlling for resource quality diet and host clone. Since we only sampled microcystin-LR concentration in a subset of the beakers during parasite exposure, we averaged together all subsampled replicates from the same treatment and applied the average to each replicate within the treatment. Since there were few fungal infected Mid37s, we were unable to include host clone in the model.

In both the fungus and bacterium experiments, microcystin-LR concentration did not impact the ultimate mature spore yield (Table S1). Resource quality was not a significant predictor of fungus mature spore yield when microcystin-LR concentration was included in the model, in contrast to the results discussed in the main text where hosts fed high-quality diets produced more spores. Host clone was the only strong predictor of bacterial spore yield, with Mid37s generally producing more mature spores than Standards.

**Table S1.** Type II analysis of deviance tables for the effect of microcystin-LR concentration on mature fungal and bacterial spore yield while controlling for diet and host clone effects. The fungus model could not include a term for host clone due to too few Mid37s infected in some diet treatments. Significant p-values are bolded.

| Response variable | Main Factors | Chi square | df | P value |
|---|---|---|---|---|
| *Fungus Mature Spore Yield* | **Avg microcystin concentration** | 2.41 | 1 | 0.12 |
| | **Diet** | 5.81 | 3 | 0.12 |
| *Bacterium Mature Spore Yield* | **Avg microcystin concentration** | 0.75 | 1 | 0.39 |
| | **Diet** | 5.62 | 3 | 0.13 |
| | **Clone** | 11.16 | 1 | **0.0008** |

*S1.6 Results for feeding rate and infection prevalence*

We conducted a test of whether feeding rate predicted the likelihood of infection for both the fungus and bacterium experiments. We used a GLMM with a binomial distribution and logit link function with binary infection presence or absence for each individual as the response variable. We tested the main factors of host clone and feeding rate during parasite exposure, and their interaction with random effects of resource quality diet and block.

In the fungus experiment, feeding rate was not correlated with infection prevalence ($\chi^2_{\text{feeding rate}}$ = 0.18, df = 1, p = 0.67). There was a marginal, but not significant effect of host clone, where Standards were more likely to be infected than Mid37s ($\chi^2_{\text{clone}}$ = 3.16, df = 1, p =0.075), and the interaction between feeding rate and host clone was not significant (Figure S3a; $\chi^2_{\text{clone×feeding rate}}$ = 0.50, df = 1, p =0.48). In the bacterium experiment, neither host clone nor feeding rate significantly impacted bacterial infection prevalence ($\chi^2_{\text{clone}}$ = 1.01, df = 1, p =0.31; $\chi^2_{\text{feeding rate}}$ = 0.07, df = 1, p = 0.79). However, there was a marginal, but not significant, interaction between host clone and feeding rate (Figure S3b; $\chi^2_{\text{clone×feeding rate}}$ = 2.95, df = 1, p =0.086). Mid37s that fed at higher rates while consuming the higher quality S or SM diets were more likely to become infected than Mid37s that fed at substantially lower rates on the M and M+ diets. Meanwhile, Standards did not have any strong association between feeding rate and infection prevalence.

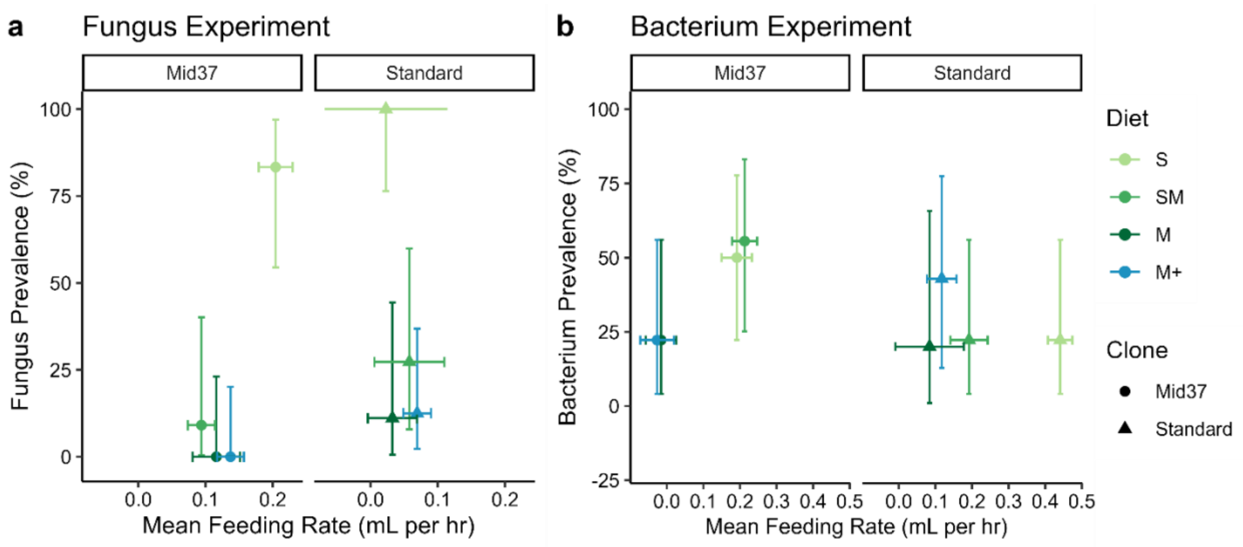


**Figure S3.** a) Fungus prevalence and b) bacterium prevalence were not associated with feeding rate. The shaded confidence intervals for a) were very large and therefore removed to maintain consistent y-axes between the two panels. Each point represents the prevalence for a given host clone and resource quality diet based on that treatment's average feeding rate among animals inoculated with parasites. The vertical error bars are the 95% confidence intervals on the parasite prevalence, and the horizontal error bars are the standard error on the average feeding rate.

*S1.7 Analysis and results for relative feeding rate and body size during parasite exposure*

We tested how body size-corrected feeding rate (i.e., relative feeding rate) during parasite exposure differed based on resource quality diets, host clone, infection status, and their interactions using a linear mixed effects model and block as a random effect. The relative feeding rate allows us to evaluate how much relative energetic intake the hosts are ingesting while accounting for potential differences in host body size, since larger *Daphnia* feed at higher rates than smaller *Daphnia* (Hall et al. 2007). We also tested whether there were differences in body size during the exposure period among all treatment combinations and their interactions using a linear mixed effects model and block as a random effect. Body size was log transformed to improve the fit with a normal distribution. For the bacterium experiment model, three low outlier points were removed from the body size data set to improve the normality of the model residuals to fit a Gaussian distribution (M+ Uninfected Mid37 Rep 9; M+ Bacterium Mid37 Rep 2; SM Bacterium Standard Rep 7). For both the relative feeding rate and body size models, we were able to include all two-way and three-way interactions for the bacterium experiment, but only the two-way interactions for the fungus experiment. We then followed up all models with a post-hoc test to compare the effect of diet for each host clone and infection status combination, and the effect of infection status for each clone with the diet treatments averaged together. For the body size models, we also conducted a post-hoc test to assess the differences in size during exposure between the two host clones.

Overall, the relative body-size corrected feeding rate results were similar to the non-corrected feeding rate results presented in the main text. In the fungus experiment, we found that the change in relative feeding rate with decreasing resource quality differed based on the host's infection status ($\chi^2_{diet\times infection} = 31.63$, df = 6, $p < 0.001$, Table S2). For uninfected (and uninoculated) hosts, reducing resource quality was correlated with reduced feeding rate. However, hosts that were inoculated with the fungus and that either became infected or resisted infection did not differ in their relative feeding rates based on resource quality diet, indicating that animals in the parasite inoculation treatment had similar exposures to the fungus spores in the water. Additionally, there were no significant differences between the three infection statuses within each host clone (Table S4). However, the fungus exposed but uninfected and infected Mid37's had higher relative feeding rates than Standards, while there was no difference between host clones when they were uninfected ($\chi^2_{infection\times clone} = 6.5$, df = 2, $p = 0.039$).

In the bacterium experiment, both host clones increased their relative feeding rates in the presence of the bacterial spores in the water relative to the uninfected, uninoculated hosts ($\chi^2_{infection} = 15.1$, df = 2, $p = 0.0005$, Table S2). Additionally, relative feeding rate declined in response to decreasing resource quality ($\chi^2_{diet} = 59.75$, df = 3, $p < 0.0001$). Additionally, in Standards, relative feeding rate was higher for the exposed-but-uninfected individuals when exposed to the bacterium relative to the uninfected, uninoculated hosts, but Mid37 relative feeding rate did not change based on infection status (Table S4). There were slight differences in relative feeding rate between the two host clones ($\chi^2_{clone} = 4.44$, df = 1, $p = 0.035$), but there were no significant interactions.

Host body size was impacted by both host clone and declining resource quality (Figure 4, Figure S4, Table S2). In both the fungus and bacterium experiments, Standards were larger than Mid37s during parasite exposure ($p_{fungus} < 0.0001$, $p_{bacterium} = < 0.0001$; Table S2) by, on average, 135 and 38 µm, respectively. Animals fed the highest quality S diet were consistently larger than animals fed the poorer quality diets (SM, M, and M+), and the differences based on diet tended to be larger for Standards compared to Mid37s. In the fungus experiment, there were no differences in body size at exposure between the three infection statuses for either clone (Table S4). In the bacterium experiment, Mid37 size did not differ based on infection status. However, Standard body size during bacterial exposure was significantly smaller for animals that ultimately became infected compared to either uninfected or bacterium-exposed but uninfected animals. Focusing just on the animals that were inoculated with the parasite, this suggests that smaller Standards may be more susceptible to the bacterium, with animals that were larger being more likely to resist infection.

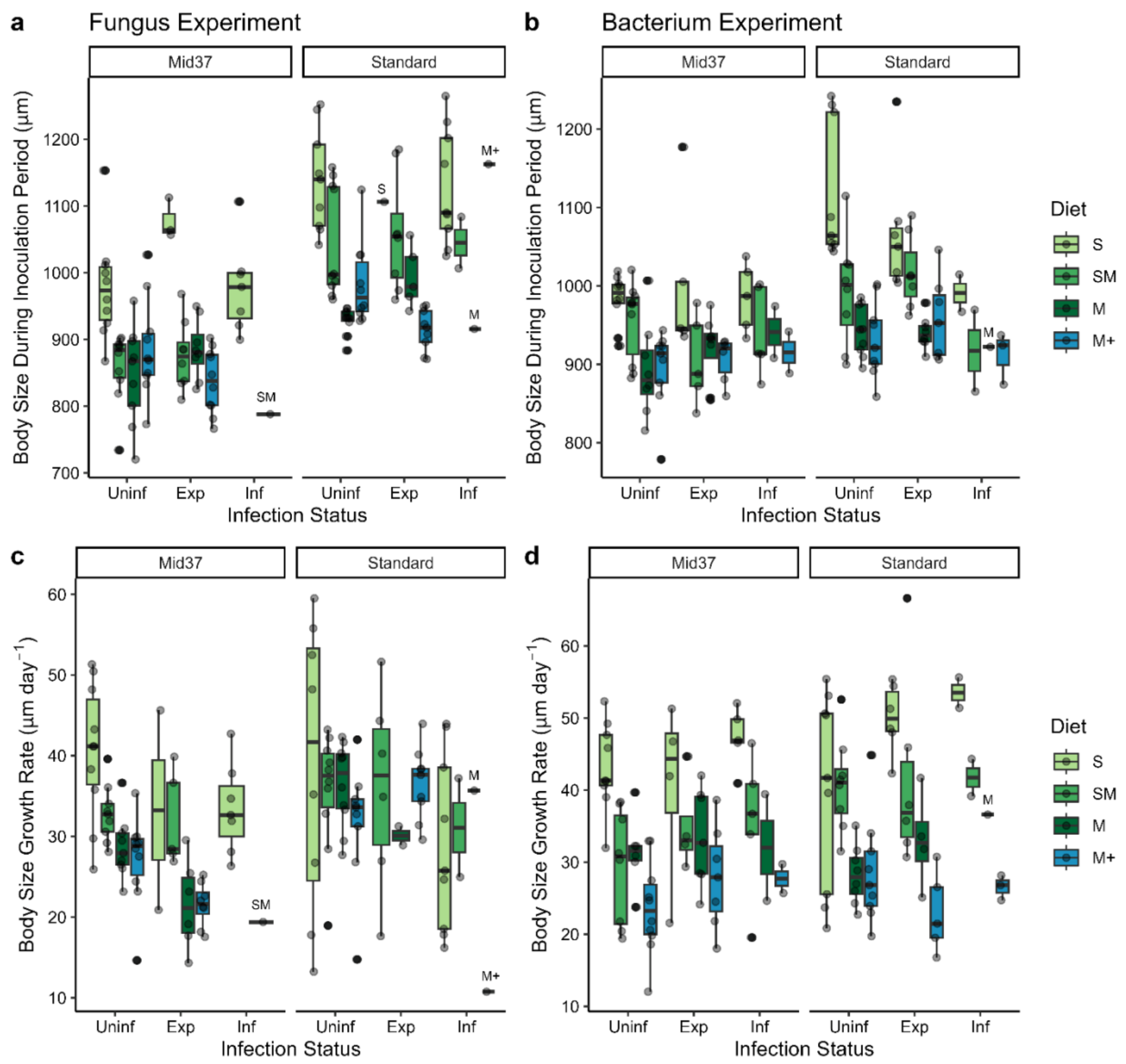


**Figure S4.** (a, b) Body size during the inoculation period (experiment days 7 and 8), and (c, d) body size growth rate from parasite exposure to the following week for the fungus experiment (a,c) and bacterium experiment (b,d), respectively. Infection status: Uninf = Uninfected (and uninoculated), Exp = Exposed but uninfected, Inf = Infected.

*S1.8 Analysis and results for host growth and metabolic rate*

We tested the effects of resource quality, host clone, and infection status on host growth rate per day from during parasite exposure to the following week for both experiments and host metabolic rate during parasite exposure. Growth rate and metabolic rate were modeled with LMMs that included diet, host clone, infection status, all two-way interactions, and Block as a random effect. The models for the bacterium experiment included all two-way and three-way interactions, while the fungus experiment models only had sufficient power to include the two-way interactions. Animals that died prior to experimental day 15, when the second body size measurements were taken, were not included in the growth rate analysis because we could not calculate a growth rate for those individuals. For the metabolic rate analysis, we removed two outliers from the fungus data set to improve the fit of the model residuals. We followed up these models with two Tukey-adjusted post-hoc tests to assess growth rate and metabolic rate differences based on diet among host clone and infection status combinations, and differences in growth rate and metabolic rate among the three infection statuses for each clone.

Host body size growth rate per day declined with reducing resource quality in both experiments and differed among host clones, and infection status also impacted growth rate in the fungus experiment (Table S2). In general, the growth rate of Mid37 was sensitive to decreasing resource quality, while Standards maintained similar high growth rates regardless of diet (Figure S4e; $\chi^2_{diet} = 23.9$, df = 3, $p < 0.001$, $\chi^2_{clone} = 17.4$, df = 1, $p < 0.001$, $\chi^2_{diet \times clone} = 6.2$, df = 3, p = 0.10). Growth rate also differed based on fungus infection status ($\chi^2_{infection} = 17.4$, df = 2, $p < 0.001$). Fungus-exposed-but-uninfected Mid37s grew 5.9 µm per day less than uninfected Mid37s, but neither the uninfected nor exposed Mid37s differed significantly from the infected animals. For Standards, there was no significant difference between uninfected and fungus-exposed-but-uninfected hosts, but infected Standards grew 9.4 µm per day less than uninfected Standards.

In the bacterium experiment, all hosts had reduced growth rates as resource quality declined regardless of host clone or infection status (Figure S4f; $\chi^2_{diet} = 99.0$, df = 3, $p < 0.001$). Host clones marginally differed in their growth rates, with Standards growing slightly faster than Mid37s ($\chi^2_{clone} = 3.9$, df = 1, p = 0.049). There was a marginal effect of infection on growth rate ($\chi^2_{infection} = 5.9$, df = 2, p = 0.051), but there were no significant differences among infection statuses (Table S4).

Host metabolic rate generally declined with poorer quality diets and increased with exposure to the fungus for one host clone, but metabolic rate was less affected by exposure to the bacterium. In the fungus experiment, metabolic rate during the parasite exposure period was lower for animals fed poorer quality resource diets and differed based on host clone (Figure S5a, Table S2, S4; $\chi^2_{diet} = 22.1$, df = 3, $p < 0.0001$, $\chi^2_{clone} = 5.7$, df = 1, p = 0.017). There was a marginal effect of infection on metabolic rate during exposure ($\chi^2_{infection} = 5.2$, df = 2, p = 0.075); this was driven by fungus-exposed-but-uninfected Mid37s having average metabolic rates that were about double those of uninoculated, uninfected Mid37s (p = 0.026; Table S4). For the bacterium experiment, metabolic rate was most affected by diet (Figure S5b, Table S2, S4; $\chi^2_{diet}$ =13.5, df = 3, p = 0.0037). The Standards also had slightly higher metabolic rate than Mid37s ($\chi^2_{clone} = 4.0$, df = 1, p = 0.045). There were no differences in metabolic rate based on their ultimate infection status (Table S4).

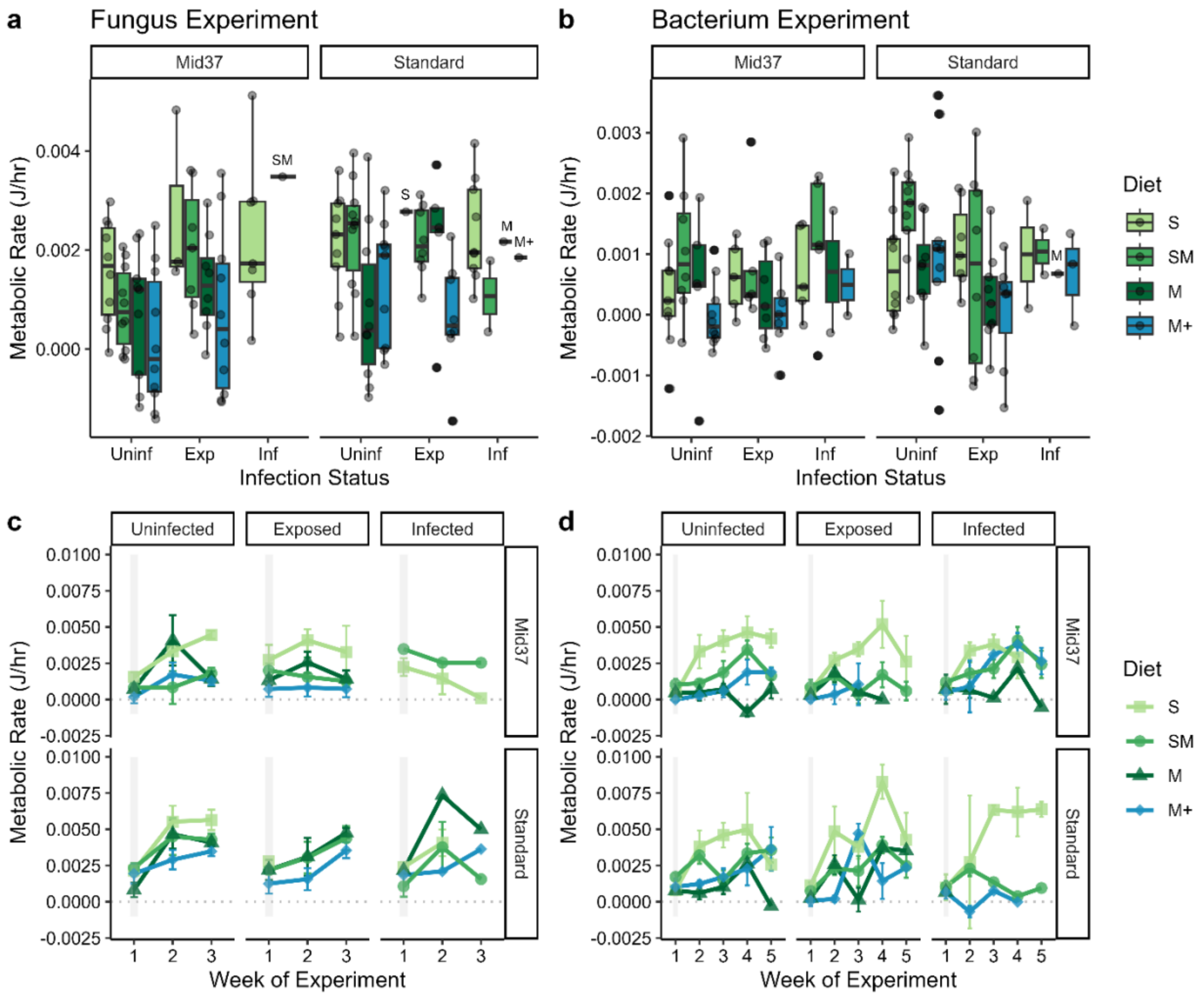


**Figure S5.** Metabolic rate during parasite exposure (a,b) and average metabolic rate per treatment during each week of the experiment with standard error (SE) confidence intervals (c,d) for the fungus (a,c) and bacterium experiments (b,d). In panels (c) and (d), points without error bars have a single replicate for that treatment combination and time point. The gray shaded box in (c) and (d) indicate the duration of parasite exposure in the first week of the experiment. Infection status: Uninf = Uninfected, Exp = Exposed but uninfected, Inf = Infected.

**Table S2.** Type II analysis of deviance tables for response variables for the fungus and bacterium experiments, respectively, that are not presented in the main text: feeding rate, relative feeding rate and body size during inoculation period, host total fecundity, body size growth rate from parasite exposure to the following week, and metabolic rate during parasite exposure. Significant p-values are bolded.

| **Response Variable** | **Predictor Variables** | **Fungus Experiment** | | | **Bacterium Experiment** | | |
|---|---|---|---|---|---|---|---|
| | | **Chi-square** | **DF** | **P-value** | **Chi-square** | **DF** | **P-value** |
| ***Relative Body Size Corrected Feeding Rate*** | Diet | 19.02 | 3 | **0.00027** | 59.75 | 3 | **<0.0001** |
| | Infection | 7.35 | 2 | **0.025** | 15.05 | 2 | **0.0005** |
| | Clone | 16.39 | 1 | **0.00005** | 4.44 | 1 | **0.035** |
| | Diet × Infection | 31.63 | 6 | **0.00002** | 4.30 | 6 | 0.64 |
| | Diet × Clone | 3.70 | 3 | 0.30 | 7.01 | 3 | 0.072 |
| | Infection × Clone | 6.49 | 2 | **0.039** | 0.78 | 2 | 0.68 |
| | Diet × Infection × Clone | -- | -- | -- | 5.17 | 6 | 0.52 |
| ***Body Size during Inoculation Period*** | Diet | 129.75 | 3 | **<0.0001** | 101.00 | 3 | **<0.0001** |
| | Infection | 0.55 | 2 | 0.76 | 2.12 | 2 | 0.35 |
| | Clone | 180.78 | 1 | **<0.0001** | 33.33 | 1 | **<0.0001** |
| | Diet × Infection | 28.94 | 6 | **<0.0001** | 5.35 | 6 | 0.50 |
| | Diet × Clone | 14.10 | 3 | **0.0028** | 5.67 | 3 | 0.13 |
| | Infection × Clone | 0.56 | 2 | 0.76 | 7.92 | 2 | **0.019** |
| | Diet × Infection × Clone | -- | -- | -- | 7.28 | 6 | 0.30 |
| ***Body Size Growth Rate*** | Diet | 23.92 | 3 | **<0.0001** | 99.03 | 3 | **<0.0001** |
| | Infection | 17.40 | 2 | **0.00017** | 5.93 | 2 | 0.051 |
| | Clone | 12.11 | 1 | **0.0005** | 3.87 | 1 | **0.049** |
| | Diet × Infection | 5.27 | 6 | 0.51 | 2.47 | 6 | 0.87 |
| | Diet × Clone | 6.22 | 3 | 0.102 | 7.01 | 3 | 0.071 |
| | Infection × Clone | 2.33 | 2 | 0.31 | 0.05 | 2 | 0.98 |
| | Diet × Infection × Clone | -- | -- | -- | 8.87 | 6 | 0.18 |
| ***Metabolic Rate*** | Diet | 22.09 | 3 | **<0.0001** | 13.49 | 3 | **0.0037** |
| | Infection | 5.19 | 2 | 0.075 | 2.49 | 2 | 0.29 |
| | Clone | 5.74 | 1 | **0.017** | 4.01 | 1 | **0.045** |
| | Diet × Infection | 4.01 | 6 | 0.68 | 4.21 | 6 | 0.65 |
| | Diet × Clone | 0.36 | 3 | 0.95 | 0.96 | 3 | 0.81 |
| | Infection × Clone | 3.29 | 2 | 0.19 | 2.88 | 2 | 0.24 |
| | Diet × Infection × Clone | -- | -- | -- | 1.42 | 6 | 0.96 |

**Table S3.** Contrasts for feeding rate in the fungus experiment among different resource quality levels conditional on the host clone and infection status. 'Uninfected' indicates animals that were uninoculated, 'Exposed' indicates animals that were exposed-but-uninfected, and 'Infected' indicates animals that became infected. P-value adjusted using the Tukey method for comparing a family of 4 estimates among the diet treatments within each host clone and infection status. Significant contrasts are bolded.

| Infection | Clone | Diet Contrast | Estimate | SE | df | t ratio | p-value |
|---|---|---|---|---|---|---|---|
| **Uninfected** | Mid37 | S – SM | -0.07 | 0.06 | 130 | -1.2 | 0.63 |
| | | S – M | 0.13 | 0.06 | 130 | 2.2 | 0.14 |
| | | **S – M+** | **0.19** | **0.06** | **130** | **3.4** | **0.005** |
| | | **SM – M** | **0.19** | **0.06** | **130** | **3.5** | **0.004** |
| | | **SM – M+** | **0.25** | **0.05** | **130** | **4.8** | **<0.0001** |
| | | M – M+ | 0.06 | 0.06 | 130 | 1.1 | 0.67 |
| **Exposed** | Mid37 | S – SM | -0.08 | 0.08 | 130 | -1.0 | 0.76 |
| | | S – M | -0.07 | 0.08 | 130 | -0.9 | 0.82 |
| | | S – M+ | -0.10 | 0.08 | 130 | -1.3 | 0.60 |
| | | SM – M | 0.01 | 0.06 | 130 | 0.1 | 1.00 |
| | | SM – M+ | -0.02 | 0.06 | 130 | -0.3 | 0.99 |
| | | M – M+ | -0.03 | 0.06 | 130 | -0.5 | 0.97 |
| **Infected** | Mid37 | S – SM | -0.07 | 0.10 | 130 | -0.7 | 0.892 |
| | | S – M | 0.01 | 0.16 | 130 | 0.1 | 1.000 |
| | | S – M+ | -0.11 | 0.16 | 130 | -0.7 | 0.911 |
| | | SM – M | 0.08 | 0.17 | 130 | 0.5 | 0.968 |
| | | SM – M+ | -0.04 | 0.17 | 130 | -0.2 | 0.996 |
| | | M – M+ | -0.11 | 0.20 | 130 | -0.6 | 0.944 |
| **Uninfected** | Standard | S – SM | -0.001 | 0.06 | 130 | 0.0 | 1 |
| | | S – M | 0.15 | 0.06 | 130 | 2.6 | 0.051 |
| | | **S – M+** | **0.27** | **0.06** | **130** | **4.8** | **<0.0001** |
| | | **SM – M** | **0.15** | **0.05** | **130** | **2.8** | **0.030** |
| | | **SM – M+** | **0.27** | **0.05** | **130** | **5.2** | **<0.0001** |
| | | M – M+ | 0.12 | 0.05 | 130 | 2.3 | 0.108 |
| **Exposed** | Standard | S – SM | -0.01 | 0.09 | 130 | -0.2 | 0.999 |
| | | S – M | -0.05 | 0.09 | 130 | -0.5 | 0.959 |
| | | S – M+ | -0.01 | 0.09 | 130 | -0.1 | 0.999 |
| | | SM – M | -0.03 | 0.06 | 130 | -0.5 | 0.955 |
| | | SM – M+ | 0.001 | 0.06 | 130 | 0.0 | 1.000 |
| | | M – M+ | 0.03 | 0.06 | 130 | 0.5 | 0.949 |
| **Infected** | Standard | S – SM | -0.004 | 0.09 | 130 | -0.1 | 1.000 |
| | | S – M | 0.03 | 0.14 | 130 | 0.2 | 0.995 |
| | | S – M+ | -0.02 | 0.14 | 130 | -0.1 | 0.999 |
| | | SM – M | 0.04 | 0.16 | 130 | 0.2 | 0.995 |
| | | SM – M+ | -0.02 | 0.16 | 130 | -0.1 | 1.000 |
| | | M – M+ | -0.05 | 0.19 | 130 | -0.3 | 0.992 |

**Table S4.** Contrasts for feeding rate, relative feeding rate, body size growth rate, metabolic rate, total lifetime fecundity and body size during parasite exposure in the fungus and bacterium experiments among different infection statuses conditional on the host clone and averaged over all resource quality diet levels. ‘Uninfected’ indicates animals that were uninoculated, ‘Exposed’ indicates animals that were exposed-but-uninfected, and ‘Infected’ indicates animals that became infected. P-value adjusted using the Tukey method for comparing a family of 3 estimates.

| Response Variable | Clone | Infection Contrast | Fungus Experiment | | | | | Bacterium Experiment | | | | |
|---|---|---|---|---|---|---|---|---|---|---|---|---|
| | | | **Estimate** | **SE** | **df** | **t ratio** | **p-value** | **Estimate** | **SE** | **df** | **t ratio** | **p-value** |
| ***Feeding Rate/ Parasite Exposure*** | Mid37 | Uninfected – Exposed | -0.05 | 0.04 | 130 | -1.5 | 0.31 | -0.08 | 0.04 | 126 | -1.86 | 0.16 |
| | | Uninfected – Infected | -0.17 | 0.08 | 130 | -2.0 | 0.12 | -0.08 | 0.05 | 125 | -1.57 | 0.26 |
| | | Exposed – Infected | -0.11 | 0.09 | 130 | -1.3 | 0.40 | -0.01 | 0.06 | 126 | -0.09 | 1.00 |
| | Standard | Uninfected – Exposed | -0.01 | 0.04 | 130 | -0.3 | 0.94 | **-0.14** | **0.04** | **126** | **-3.40** | **0.0026** |
| | | Uninfected – Infected | 0.02 | 0.06 | 130 | 0.4 | 0.93 | -0.07 | 0.07 | 126 | -0.97 | 0.60 |
| | | Exposed – Infected | 0.03 | 0.06 | 130 | 0.5 | 0.85 | 0.07 | 0.07 | 125 | 1.01 | 0.57 |
| ***Relative Body Size Corrected Feeding Rate*** | Mid37 | Uninfected – Exposed | -0.07 | 0.04 | 130 | -1.9 | 0.16 | -0.10 | 0.05 | 126 | -1.93 | 0.13 |
| | | Uninfected – Infected | -0.19 | 0.09 | 130 | -2.1 | 0.09 | -0.09 | 0.06 | 125 | -1.50 | 0.29 |
| | | Exposed – Infected | -0.12 | 0.09 | 130 | -1.3 | 0.41 | 0.002 | 0.07 | 126 | 0.03 | 1.00 |
| | Standard | Uninfected – Exposed | -0.02 | 0.04 | 130 | -0.5 | 0.87 | **-0.15** | **0.05** | **126** | **-3.15** | **0.006** |
| | | Uninfected – Infected | 0.02 | 0.06 | 130 | 0.4 | 0.93 | -0.08 | 0.08 | 126 | -1.00 | 0.58 |
| | | Exposed – Infected | 0.04 | 0.07 | 130 | 0.6 | 0.80 | 0.07 | 0.08 | 125 | 0.83 | 0.69 |
| ***Body Size Growth Rate*** | Mid37 | Uninfected – Exposed | **5.84** | **2.26** | **115** | **2.6** | **0.030** | -2.24 | 2.25 | 108 | -0.99 | 0.58 |
| | | Uninfected – Infected | 9.55 | 4.81 | 115 | 2.0 | 0.12 | -3.69 | 2.73 | 107 | -1.35 | 0.37 |
| | | Exposed – Infected | 3.71 | 5.04 | 115 | 0.7 | 0.74 | -1.45 | 2.95 | 108 | -0.49 | 0.88 |
| | Standard | Uninfected – Exposed | 0.73 | 2.93 | 115 | 0.3 | 0.97 | -2.46 | 2.23 | 108 | -1.11 | 0.51 |
| | | Uninfected – Infected | **9.39** | **3.29** | **115** | **2.9** | **0.014** | -5.16 | 3.37 | 108 | -1.53 | 0.28 |
| | | Exposed – Infected | 8.66 | 4.09 | 115 | 2.1 | 0.091 | -2.69 | 3.61 | 107 | -0.75 | 0.74 |
| ***Metabolic Rate*** | Mid37 | Uninfected / Exposed | **-0.0009** | **0.0003** | **129** | **-2.6** | **0.026** | -0.00003 | 0.0003 | 120 | -0.10 | 0.99 |
| | | Uninfected / Infected | -0.0015 | 0.0008 | 129 | -1.9 | 0.16 | -0.0003 | 0.0003 | 120 | -0.89 | 0.65 |
| | | Exposed / Infected | -0.0006 | 0.0008 | 129 | -0.7 | 0.75 | -0.0003 | 0.0004 | 120 | -0.77 | 0.72 |
| | Standard | Uninfected / Exposed | -0.0003 | 0.0004 | 130 | -0.7 | 0.75 | 0.0005 | 0.0002 | 120 | 2.13 | 0.09 |
| | | Uninfected / Infected | -0.0004 | 0.0005 | 129 | -0.7 | 0.79 | 0.0003 | 0.0004 | 120 | 0.77 | 0.72 |
| | | Exposed / Infected | -0.0001 | 0.0006 | 130 | -0.1 | 0.99 | -0.0002 | 0.0004 | 121 | -0.47 | 0.89 |
| | | | **Ratio** | **SE** | **df** | **z ratio** | **p-value** | **Ratio** | **SE** | **df** | **z ratio** | **p-value** |
| ***Total Fecundity*** | Mid37 | Uninfected / Exposed | 1.01 | 0.16 | Inf | 0.05 | 0.999 | **3.25** | **1.00** | **Inf** | **3.43** | **0.0017** |
| | | Uninfected / Infected | **2.69** | **0.81** | **Inf** | **3.3** | **0.0028** | 360.91 | 399504.0 | Inf | 0.005 | 1 |
| | | Exposed / Infected | **2.67** | **0.87** | **Inf** | **3.0** | **0.0073** | 110.96 | 122829.0 | Inf | 0.004 | 1 |

| | | | | | | | | | | | | |
|---|---|---|---|---|---|---|---|---|---|---|---|---|
| | Standard | Uninfected / Exposed | **1.61** | **0.22** | **Inf** | **3.4** | **0.0017** | 0.82 | 0.0000 | Inf | -0.969 | 0.60 |
| | | Uninfected / Infected | **1.97** | **0.34** | **Inf** | **4.0** | **0.0002** | 935.40 | 25678879.0 | Inf | 0 | 1 |
| | | Exposed / Infected | 1.23 | 0.25 | Inf | 1.0 | 0.57 | 1141.74 | 31343593.0 | Inf | 0 | 1 |
| ***Body size during parasite exposure*** | Mid37 | Uninfected – Exposed | 0.98 | 0.02 | 132 | -1.53 | 0.28 | 1.00 | 0.02 | 121 | -0.08 | 0.997 |
| | | Uninfected – Infected | 0.99 | 0.04 | 132 | -0.32 | 0.94 | 0.99 | 0.02 | 122 | -0.83 | 0.69 |
| | | Exposed – Infected | 1.01 | 0.04 | 132 | 0.30 | 0.95 | 0.99 | 0.02 | 123 | -0.71 | 0.76 |
| | Standard | Uninfected – Exposed | 0.99 | 0.02 | 132 | -0.73 | 0.75 | 1.00 | 0.01 | 122 | -0.10 | 0.99 |
| | | Uninfected – Infected | 0.97 | 0.03 | 132 | -1.16 | 0.48 | **1.06** | **0.03** | **123** | **2.50** | **0.037** |
| | | Exposed – Infected | 0.98 | 0.03 | 132 | -0.60 | 0.82 | **1.06** | **0.03** | **122** | **2.49** | **0.038** |

**Table S5.** Contrasts for feeding rate in the bacterium experiment among different resource quality levels conditional on the host clone and infection status. 'Uninfected' indicates animals that were uninoculated, 'Exposed' indicates animals that were exposed-but-uninfected, and 'Infected' indicates animals that became infected. P-value adjusted using the Tukey method for comparing a family of 4 estimates among the diet treatments within each host clone and infection status. Significant contrasts are bolded.

| Infection | Clone | Diet Contrast | Estimate | SE | df | t ratio | p-value |
|---|---|---|---|---|---|---|---|
| **Uninfected** | Mid37 | S – SM | -0.05 | 0.07 | 125 | -0.68 | 0.91 |
| | | **S – M** | **0.21** | **0.08** | **125** | **2.81** | **0.029** |
| | | S – M+ | 0.03 | 0.07 | 125 | 0.49 | 0.96 |
| | | **SM – M** | **0.26** | **0.08** | **125** | **3.45** | **0.004** |
| | | SM – M+ | 0.08 | 0.07 | 125 | 1.17 | 0.65 |
| | | M – M+ | -0.18 | 0.08 | 125 | -2.36 | 0.091 |
| **Exposed** | Mid37 | S – SM | -0.03 | 0.10 | 125 | -0.30 | 0.99 |
| | | S – M | 0.18 | 0.09 | 125 | 1.91 | 0.23 |
| | | S – M+ | 0.23 | 0.09 | 126 | 2.48 | 0.069 |
| | | SM – M | 0.21 | 0.09 | 125 | 2.23 | 0.12 |
| | | **SM – M+** | **0.26** | **0.09** | **126** | **2.79** | **0.031** |
| | | M – M+ | 0.05 | 0.08 | 126 | 0.64 | 0.92 |
| **Infected** | Mid37 | S – SM | -0.03 | 0.10 | 125 | -0.34 | 0.99 |
| | | S – M | 0.27 | 0.14 | 120 | 1.91 | 0.23 |
| | | S – M+ | 0.16 | 0.13 | 126 | 1.19 | 0.64 |
| | | SM – M | 0.30 | 0.14 | 120 | 2.16 | 0.14 |
| | | SM – M+ | 0.19 | 0.13 | 126 | 1.44 | 0.48 |
| | | M – M+ | -0.11 | 0.16 | 126 | -0.68 | 0.91 |
| **Uninfected** | Standard | S – SM | 0.12 | 0.08 | 125 | 1.58 | 0.39 |
| | | **S – M** | **0.40** | **0.07** | **125** | **5.33** | **<.0001** |
| | | **S – M+** | **0.31** | **0.08** | **125** | **4.14** | **0.0004** |
| | | **SM – M** | **0.28** | **0.08** | **125** | **3.58** | **0.0028** |
| | | SM – M+ | 0.19 | 0.08 | 126 | 2.42 | 0.079 |
| | | M – M+ | -0.09 | 0.08 | 125 | -1.17 | 0.65 |
| **Exposed** | Standard | **S – SM** | **0.23** | **0.08** | **126** | **2.80** | **0.030** |
| | | **S – M** | **0.37** | **0.08** | **125** | **4.45** | **0.0001** |
| | | **S – M+** | **0.31** | **0.09** | **125** | **3.63** | **0.0023** |
| | | SM – M | 0.13 | 0.08 | 125 | 1.69 | 0.34 |
| | | SM – M+ | 0.08 | 0.08 | 125 | 0.95 | 0.78 |
| | | M – M+ | -0.06 | 0.08 | 125 | -0.69 | 0.90 |
| **Infected** | Standard | S – SM | 0.38 | 0.16 | 126 | 2.35 | 0.093 |
| | | **S – M** | **0.62** | **0.20** | **126** | **3.14** | **0.011** |
| | | S – M+ | 0.38 | 0.15 | 125 | 2.59 | 0.052 |
| | | SM – M | 0.24 | 0.19 | 125 | 1.23 | 0.61 |
| | | SM – M+ | 0.00 | 0.15 | 126 | -0.02 | 1 |
| | | M – M+ | -0.24 | 0.18 | 126 | -1.31 | 0.56 |

**Table S6.** Contrasts for total fecundity in the fungus experiment among different resource quality levels conditional on the host clone and infection status. 'Uninfected' indicates animals that were uninoculated, 'Exposed' indicates animals that were exposed-but-uninfected, and 'Infected' indicates animals that became infected. P-value adjusted using the Tukey method for comparing a family of 4 estimates among the diet treatments within each host clone and infection status. Significant contrasts are bolded.

| Infection | Clone | Diet Contrast | Ratio | SE | df | z ratio | p-value |
|---|---|---|---|---|---|---|---|
| **Uninfected** | Mid37 | **S – SM** | **3.60** | **0.83** | **Inf** | **5.58** | **<0.0001** |
| | | **S – M** | **4.64** | **1.15** | **Inf** | **6.20** | **<0.0001** |
| | | **S – M+** | **4.41** | **1.03** | **Inf** | **6.38** | **<0.0001** |
| | | SM – M | 1.29 | 0.33 | Inf | 0.98 | 0.87 |
| | | SM – M+ | 1.23 | 0.30 | Inf | 0.82 | 0.91 |
| | | M – M+ | 0.95 | 0.25 | Inf | -0.19 | 1.0 |
| **Exposed** | Mid37 | **S – SM** | **2.48** | **0.85** | **Inf** | **2.64** | **0.041** |
| | | **S – M** | **3.50** | **1.22** | **Inf** | **3.58** | **0.002** |
| | | **S – M+** | **3.92** | **1.36** | **Inf** | **3.93** | **0.001** |
| | | SM – M | 1.41 | 0.33 | Inf | 1.46 | 0.46 |
| | | SM – M+ | 1.58 | 0.36 | Inf | 1.98 | 0.19 |
| | | M – M+ | 1.12 | 0.27 | Inf | 0.47 | 0.97 |
| **Infected** | Mid37 | **S – SM** | **4.52** | **1.73** | **Inf** | **3.95** | **0.0005** |
| | | S – M | 2.81 | 1.45 | Inf | 2.01 | 0.19 |
| | | **S – M+** | **5.06** | **2.46** | **Inf** | **3.35** | **0.0046** |
| | | SM – M | 0.62 | 0.35 | Inf | -0.85 | 0.83 |
| | | SM – M+ | 1.12 | 0.59 | Inf | 0.21 | 1.00 |
| | | M – M+ | 1.80 | 1.14 | Inf | 0.94 | 0.79 |
| **Uninfected** | Standard | S – SM | 0.99 | 0.14 | Inf | -0.07 | 1.00 |
| | | S – M | 1.41 | 0.22 | Inf | 2.20 | 0.12 |
| | | S – M+ | 1.45 | 0.23 | Inf | 2.35 | 0.087 |
| | | SM – M | 1.42 | 0.20 | Inf | 2.52 | 0.056 |
| | | **SM – M+** | **1.47** | **0.21** | **Inf** | **2.69** | **0.036** |
| | | M – M+ | 1.03 | 0.16 | Inf | 0.22 | 1.00 |
| **Exposed** | Standard | S – SM | 0.68 | 0.27 | Inf | -0.96 | 0.77 |
| | | S – M | 1.06 | 0.44 | Inf | 0.14 | 1.00 |
| | | S – M+ | 1.29 | 0.52 | Inf | 0.63 | 0.92 |
| | | SM – M | 1.55 | 0.30 | Inf | 2.28 | 0.10 |
| | | **SM – M+** | **1.89** | **0.33** | **Inf** | **3.70** | **0.001** |
| | | M – M+ | 1.22 | 0.23 | Inf | 1.04 | 0.72 |
| **Infected** | Standard | S – SM | 1.25 | 0.37 | Inf | 0.74 | 0.88 |
| | | S – M | 0.85 | 0.37 | Inf | -0.37 | 0.98 |
| | | S – M+ | 1.67 | 0.68 | Inf | 1.25 | 0.59 |
| | | SM – M | 0.68 | 0.34 | Inf | -0.77 | 0.87 |
| | | SM – M+ | 1.34 | 0.64 | Inf | 0.62 | 0.93 |
| | | M – M+ | 1.96 | 1.12 | Inf | 1.17 | 0.64 |

**Table S7.** Contrasts for total fecundity in the bacterium experiment among different resource quality levels conditional on the host clone and infection status. 'Uninfected' indicates animals that were uninoculated, 'Exposed' indicates animals that were exposed-but-uninfected, and 'Infected' indicates animals that became infected. P-value adjusted using the Tukey method for comparing a family of 4 estimates among the diet treatments within each host clone and infection status. Significant contrasts are bolded.

| Infection | Clone | Diet Contrast | Ratio | SE | df | z ratio | p-value |
|---|---|---|---|---|---|---|---|
| **Uninfected** | Mid37 | S – SM | 2.00 | 1.00 | Inf | 1.59 | 0.38 |
| | | **S – M** | **8.00** | **3.00** | **Inf** | **4.33** | **0.0001** |
| | | **S – M+** | **8.00** | **3.00** | **Inf** | **5.43** | **<0.0001** |
| | | **SM – M** | **4.00** | **2.00** | **Inf** | **2.93** | **0.018** |
| | | **SM – M+** | **5.00** | **2.00** | **Inf** | **3.75** | **0.001** |
| | | M – M+ | 1.00 | 0.00 | Inf | 0.35 | 0.99 |
| **Exposed** | Mid37 | S – SM | 3.00 | 2.00 | Inf | 1.92 | 0.22 |
| | | **S – M** | **8.00** | **5.00** | **Inf** | **3.83** | **0.0007** |
| | | **S – M+** | **90.00** | **98.00** | **Inf** | **4.12** | **0.0002** |
| | | SM – M | 3.00 | 2.00 | Inf | 1.81 | 0.27 |
| | | **SM – M+** | **32.00** | **35.00** | **Inf** | **3.10** | **0.011** |
| | | M – M+ | 11.00 | 12.00 | Inf | 2.15 | 0.14 |
| **Infected** | Mid37 | S – SM | 2.00 | 1.00 | Inf | 0.84 | 0.84 |
| | | S – M | 9.19E+07 | 4.07E+11 | Inf | 0.004 | 1 |
| | | S – M+ | 2.00 | 2.00 | Inf | 0.90 | 0.80 |
| | | SM – M | 5.34E+07 | 2.36E+11 | Inf | 0.004 | 1 |
| | | SM – M+ | 1.00 | 1.00 | Inf | 0.33 | 0.99 |
| | | M – M+ | 0.00 | 0.00 | Inf | -0.004 | 1 |
| **Uninfected** | Standard | S – SM | 1.00 | 0.00 | Inf | 0.21 | 1.00 |
| | | **S – M** | **4.00** | **1.00** | **Inf** | **3.65** | **0.0015** |
| | | S – M+ | 2.00 | 1.00 | Inf | 2.04 | 0.17 |
| | | **SM – M** | **4.00** | **1.00** | **Inf** | **3.36** | **0.0044** |
| | | SM – M+ | 2.00 | 1.00 | Inf | 1.77 | 0.29 |
| | | M – M+ | 1.00 | 0.00 | Inf | -1.66 | 0.35 |
| **Exposed** | Standard | S – SM | 1.00 | 0.00 | Inf | -0.55 | 0.95 |
| | | S – M | 3.00 | 1.00 | Inf | 2.19 | 0.13 |
| | | S – M+ | 2.00 | 1.00 | Inf | 2.06 | 0.17 |
| | | **SM – M** | **3.00** | **1.00** | **Inf** | **2.74** | **0.031** |
| | | SM – M+ | 3.00 | 1.00 | Inf | 2.55 | 0.053 |
| | | M – M+ | 1.00 | 0.00 | Inf | -0.17 | 1.00 |
| **Infected** | Standard | S – SM | 2.00 | 1.00 | Inf | 0.60 | 0.93 |
| | | S – M | 2.66E+11 | 2.93E+16 | Inf | 0.00 | 1 |
| | | **S – M+** | **16.00** | **14.00** | **Inf** | **3.38** | **0.0041** |
| | | SM – M | 1.66E+11 | 1.86E+16 | Inf | 0.00 | 1 |
| | | **SM – M+** | **10.00** | **9.00** | **Inf** | **2.81** | **0.026** |
| | | M – M+ | 0.00 | 0.00 | Inf | 0.00 | 1 |

**Table S8.** Cox proportional hazard ratios of mortality risk from the fungus and bacterium experiments based on resource quality, infection status, and host clone. 'Uninfected' indicates animals that were uninoculated, 'Exposed' indicates animals that were exposed-but-uninfected, and 'Infected' indicates animals that became infected. Hazard ratios and p-values for treatment levels that differed significantly from the reference in parentheses (i.e., S diet, uninfected, or Mid37 clone) are bolded.

| **Fungus experiment** | | | | | | | |
|---|---|---|---|---|---|---|---|
| | **Hazard Ratio** | **95% CI** | **Coefficient** | **SE** | **Z** | **Wald $\chi^2$** | **P-value** |
| *(Diet-S)* | Reference | | | | | | |
| Diet-SM | **0.24** | (0.08, 0.70) | -1.41 | 0.54 | -2.63 | 6.90 | **0.009** |
| Diet-M | **0.29** | (0.09, 0.92) | -1.24 | 0.59 | -2.11 | 4.44 | **0.035** |
| Diet-M+ | **0.27** | (0.09, 0.81) | -1.29 | 0.55 | -2.34 | 5.48 | **0.019** |
| *(Infection-Uninfected)* | Reference | | | | | | |
| Infection-Exposed | **15.09** | (3.27, 69.62) | 2.71 | 0.78 | 3.48 | 12.10 | **0.001** |
| Infection-Infected | **38.04** | (8.60, 168.35) | 3.64 | 0.76 | 4.80 | 12.99 | **<0.0001** |
| *(Clone-Mid37)* | Reference | | | | | | |
| Clone-Standard | 1.75 | (0.92, 3.33) | 0.56 | 0.33 | 1.69 | 2.86 | 0.091 |
| **Bacterium Experiment** | | | | | | | |
| *(Diet-S)* | Reference | | | | | | |
| Diet-SM | **0.33** | (0.15, 0.73) | -1.09 | 0.40 | -2.76 | 7.63 | **0.006** |
| Diet-M | **1.86** | (1.02, 3.38) | 0.62 | 0.31 | 2.04 | 4.14 | **0.042** |
| Diet-M+ | 1.46 | (0.82, 2.60) | 0.38 | 0.29 | 1.28 | 1.64 | 0.200 |
| *(Infection-Uninfected)* | Reference | | | | | | |
| Infection-Exposed | **1.67** | (1.03, 2.71) | 0.52 | 0.25 | 2.10 | 4.40 | **0.036** |
| Infection-Infected | 1.03 | (0.52, 2.06) | 0.03 | 0.35 | 0.09 | 0.01 | 0.93 |
| *(Clone-Mid37)* | Reference | | | | | | |
| Clone-Standard | 1.33 | (0.85, 2.08) | 0.28 | 0.23 | 1.24 | 1.53 | 0.22 |

**Table S9.** Cox proportional hazard ratios of mortality risk from the fungus and bacterium experiments run separately for each clone based on resource quality and infection status. 'Uninfected' indicates animals that were uninoculated, 'Exposed' indicates animals that were exposed-but-uninfected, and 'Infected' indicates animals that became infected. Hazard ratios and p-values for treatment levels that differed significantly from the reference in parentheses (i.e., S diet, uninfected) are bolded.

| **Fungus experiment: Mid37** | | | | | | | |
|---|---|---|---|---|---|---|---|
| | **Hazard Ratio** | **95% CI** | **Coefficient** | **SE** | **Z** | **Wald $\chi^2$** | **P-value** |
| *(Diet-S)* | Reference | | | | | | |
| Diet-SM | 0.46 | (0.08, 2.69) | -0.79 | 0.91 | -0.87 | 0.75 | 0.39 |
| Diet-M | 1.15 | (0.15, 8.96) | 0.14 | 1.05 | 0.14 | 0.02 | 0.89 |
| Diet-M+ | 0.24 | (0.02, 3.48) | -1.41 | 1.36 | -1.04 | 1.08 | 0.30 |
| *(Infection-Uninfected)* | Reference | | | | | | |
| Infection-Exposed | 6.09 | (0.67, 54.99) | 1.81 | 1.12 | 1.61 | 2.59 | 0.108 |
| Infection-Infected | **48.68** | (4.90, 483.29) | 3.89 | 1.17 | 3.32 | 11.01 | **0.001** |
| **Fungus experiment: Standard** | | | | | | | |
| *(Diet-S)* | Reference | | | | | | |
| Diet-SM | **0.17** | (0.04, 0.75) | -1.76 | 0.75 | -2.35 | 5.51 | **0.019** |
| Diet-M | **0.11** | (0.02, 0.71) | -2.20 | 0.95 | -2.31 | 5.35 | **0.021** |
| Diet-M+ | 0.25 | (0.06, 1.07) | -1.40 | 0.75 | -1.87 | 3.51 | 0.061 |
| *(Infection-Uninfected)* | Reference | | | | | | |
| Infection-Exposed | **28.41** | (3.17, 254.71 | 3.35 | 1.12 | 2.99 | 8.95 | **0.003** |
| Infection-Infected | **36.00** | (4.49, 288.37) | 3.58 | 1.06 | 3.38 | 11.39 | **0.001** |
| **Bacterium Experiment: Mid37** | | | | | | | |
| *(Diet-S)* | Reference | | | | | | |
| Diet-SM | **0.18** | (0.04, 0.81) | -1.74 | 0.78 | -2.24 | 5.01 | **0.025** |
| Diet-M | 1.74 | (0.72, 4.19) | 0.55 | 0.45 | 1.24 | 1.53 | 0.22 |
| Diet-M+ | 1.96 | (0.85, 4.52) | 0.67 | 0.43 | 1.58 | 2.49 | 0.11 |
| *(Infection-Uninfected)* | Reference | | | | | | |
| Infection-Exposed | **3.25** | (1.55, 6.85) | 1.18 | 0.38 | 3.11 | 9.64 | **0.002** |
| Infection-Infected | 1.18 | (0.44, 3.12) | 0.16 | 0.50 | 0.33 | 0.11 | 0.74 |
| **Bacterium Experiment: Standard** | | | | | | | |
| (Diet-S) | Reference | | | | | | |
| Diet-SM | 0.47 | (0.18, 1.20) | -0.76 | 0.48 | -1.58 | 2.50 | 0.11 |
| Diet-M | 1.87 | (0.81, 4.30) | 0.62 | 0.43 | 1.47 | 2.15 | 0.14 |
| Diet-M+ | 1.02 | (0.45, 2.30) | 0.02 | 0.42 | 0.05 | 0.00 | 0.96 |
| *(Infection-Uninfected)* | Reference | | | | | | |
| Infection-Exposed | 0.94 | (0.48, 1.85) | -0.06 | 0.34 | -0.18 | 0.03 | 0.86 |
| Infection-Infected | 1.12 | (0.41, 3.06) | 0.12 | 0.51 | 0.23 | 0.05 | 0.82 |

**Table S10. Synthesis of findings related to costs of resisting infection.** While there was substantial variation in whether there were costs seen, this variation did not appear to be mediated by diet.

| | Fungus | | Bacterium | |
|---|---|---|---|---|
| | **Mid37** | **Standard** | **Mid37** | **Standard** |
| **fecundity** | no | yes | yes | no |
| **mortality** | no | yes | yes | no |

### References Cited in Appendix S1


Alton, L. A., C. R. White, and R. S. Seymour. 2007. Effect of aerial O2 partial pressure on bimodal gas exchange and air-breathing behaviour in Trichogaster leeri. Journal of Experimental Biology **210**:2311-2319.

Bergman Filho, T. U., A. M. Soares, and S. Loureiro. 2011. Energy budget in Daphnia magna exposed to natural stressors. Environ Sci Pollut Res Int **18**:655-662.

Cameron, J. N. 1986. Appendix 2 – Solubility of O2 and CO2 at different temperatures and salinities. Pages 254-259 *in* J. Cameron, editor. Principles of physiological measurement. Academic Press, Inc.

Hall, M. D., B. L. Phillips, C. R. White, and D. J. Marshall. 2024. The hidden costs of resistance: Contrasting the energetics of successfully and unsuccessfully fighting infection. Functional Ecology **38**:714-723.

Hall, S. R., L. Sivars-Becker, C. Becker, M. A. Duffy, A. J. Tessier, and C. E. Cáceres. 2007. Eating yourself sick: transmission of disease as a function of feeding biology of hosts. Ecology Letters **10**:207-218.

Hite, J. L., A. C. Pfenning-Butterworth, R. E. Vetter, and C. E. Cressler. 2020. A high-throughput method to quantify feeding rates in aquatic organisms: A case study with Daphnia. Ecol Evol **10**:6239-6245.

Lighton, J. R. B. 2008. Flow-through respirometry using incurrent flow measurement. Pages 105-123 *in* J. R. B. Lighton, editor. Measuring Metabolic Rates: A manual for scientists. Oxford Academic.

Millard, S. P. 2013. EnvStats: An R package for environmental statistics. Springer.

Nørgaard, L. S., G. Ghedini, B. L. Phillips, and M. D. Hall. 2021. Energetic scaling across different host densities and its consequences for pathogen proliferation. Functional Ecology **35**:475-484.

Olito, C., C. R. White, D. J. Marshall, and D. R. Barneche. 2017. Estimating monotonic rates from biological data using local linear regression. Journal of Experimental Biology **220**:759-764.

Penczykowski, R. M., B. C. P. Lemanski, R. D. Sieg, S. R. Hall, J. Housley Ochs, J. Kubanek, and M. A. Duffy. 2014. Poor resource quality lowers transmission potential by changing foraging behaviour. Functional Ecology **28**:1245-1255.

Shocket, M. S., A. T. Strauss, J. L. Hite, M. Šljivar, D. J. Civitello, M. A. Duffy, C. E. Cáceres, and S. R. Hall. 2018. Warmer is sicker in a zooplankton-fungus system: a trait-driven

approach points to higher transmission via host foraging. American Naturalist **191**:435-451.
Stewart Merrill, T. E., S. Hall, L. Merrill, and C. E. Cáceres. 2019. Variation in immune defense shapes disease outcomes in laboratory and wild Daphnia. Integrative and Comparative Biology.
Strauss, A. T., J. L. Hite, D. J. Civitello, M. S. Shocket, C. E. Cáceres, and S. R. Hall. 2019. Genotypic variation in parasite avoidance behaviour and other mechanistic, nonlinear components of transmission. Proceedings of the Royal Society B: Biological Sciences **286**:20192164.
White, C. R. 2011. Allometric estimation of metabolic rates in animals. Comparative Biochemistry and Physiology Part A: Molecular & Integrative Physiology **158**:346-357.
